# Long-read single-cell genomics: resolving chimeras in multiple displacement amplification

**DOI:** 10.64898/2026.06.11.730069

**Authors:** Jamie McGowan, James Lipscombe, Estelle S. Kilias, Tom Barker, Leah Catchpole, Alex Durrant, Naomi Irish, Seanna McTaggart, Sally D. Warring, Karim Gharbi, Thomas A. Richards, Neil Hall, David Swarbreck

**Affiliations:** Earlham Institute, Norwich Research Park, Norwich, United Kingdom; School of Biomolecular and Biomedical Science, Conway Institute, University College Dublin, Dublin, Ireland; Department of Biology, University of Oxford, Oxford, United Kingdom; School of Biological Sciences, University of East Anglia, Norwich, United Kingdom

## Abstract

Multiple displacement amplification (MDA) enables whole-genome amplification from single cells, but introduces chimeric artifacts that severely compromise downstream analyses, particularly with long-read sequencing. Here, we systematically evaluate long-read PacBio HiFi sequencing of MDA amplified DNA from single cells using the model green alga *Chlamydomonas reinhardtii*. We show that MDA-derived libraries exhibit highly uneven coverage and extreme chimera rates impacting up to 70% of reads, leading to thousands of artefactual structural variants and misassemblies when assembled using algorithms designed for bulk sequencing. To overcome these challenges, we developed lrSAGA (**l**ong-**r**ead **S**ingle **A**mplified **G**enome **A**ssembly), a novel tool to assemble long-read MDA sequencing datasets. Assemblies generated using lrSAGA are more complete, more contiguous, and have 75-95% fewer misassemblies compared to conventional assembly algorithms. Although overall contiguity is limited by MDA coverage dropouts, we demonstrate that up to 68% of the *C. reinhardtii* genome can be accurately assembled from just a single haploid cell. We further validated lrSAGA using published Oxford Nanopore and PacBio HiFi data from single or half *Caenorhabditis elegans* worms, generating accurate and highly complete assemblies. Applying our approach to single protist cells isolated from environmental water samples, we performed PacBio HiFi single-cell genome sequencing of four uncultivated microbial eukaryotes: an amoeboflagellate from the *Naegleria* genus, a flagellate from the *Bodo* genus, and two deep-branching flagellates from the enigmatic CRuMs supergroup, *Collodictyon triciliatum* and *Diphylleia rotans*. From single cells, we generated high-quality draft genome assemblies estimated to be 70-84% complete, demonstrating the potential of long-read single-cell genomics to unlock genome diversity from uncultivated microbial eukaryotes.

## Introduction

In the era of “biodiversity genomics”, large-scale initiatives such as the Darwin Tree of Life Project (The Darwin Tree of Life Project Consortium 2022), the European Reference Genome Atlas (Mazzoni et al. 2023), and the Earth BioGenome Project (Blaxter et al. 2025) aim to sequence the genomes of all known eukaryotic species, with the ultimate goal of creating a genomic catalogue of global eukaryotic biodiversity. These efforts are poised to transform our understanding of ecology and evolution, and support biodiversity conservation. However, the majority of eukaryotic diversity resides among undescribed single-celled eukaryotes, collectively known as protists (Burki et al. 2020). Protists are omnipresent across nearly all ecosystems, play essential roles in biogeochemical cycles, include some of the most destructive pathogens of animals and plants, and are essential to understand both the diversification of all eukaryotes and the origin of their last common ancestor (Schoenle et al. 2025). Despite their ecological and evolutionary importance, protists tend to be overlooked in genomic research (Sibbald and Archibald 2017). Most protists are unculturable or difficult to maintain long-term under laboratory conditions. This presents a significant barrier to genome sequencing efforts. As a result, protists are vastly underrepresented in genomic databases, leaving a substantial fraction of eukaryotic diversity unexplored at the genomic level.

Recent advancements have substantially decreased the amount of input DNA required for long-read genome sequencing. For example, the Ultra-Low DNA input protocol from Pacific Biosciences (PacBio) enables PacBio HiFi sequencing from as little as 5 ng of input DNA. The recently announced Ampli-Fi protocol from PacBio is reported to enable PacBio HiFi sequencing from just 1 ng of genomic DNA. This approach relies on an alternative polymerase enzyme with reduced bias (Bein et al. 2025). Alternative approaches have lower input requirements. For example, the Picogram input Multimodal Sequencing (PiMmS) method (Laumer 2023) constructs long-read sequencing libraries from only a few hundred picograms of DNA. These developments enable genome sequencing and high quality de novo genome assembly of individual small-bodied animals. However, such amounts of input DNA are out of reach for individual single-celled organisms without culturing. For example, vegetative cells of *Chlamydomonas reinhardtii,* the model single-celled green alga used in this study, are haploid and have a nuclear genome of approximately 115 Mb (Craig et al. 2023). This equates to less than 150 femtograms of nuclear DNA per single cell.

Multiple displacement amplification (MDA) is a widely used technique for whole genome amplification (WGA) from single-cell quantities of DNA to generate and sequence single-amplified genomes (SAGs). MDA is carried out as an isothermal reaction and involves the annealing of random hexamers to denatured DNA followed by strand displacement synthesis using the phi29 polymerase (Spits et al. 2006). MDA is a desirable approach given its low error rate (1 error in 10^6^ - 10^7^ nucleotides), high yield, and relatively large product length (∼10 kb) (Dean et al. 2002). However, MDA presents several challenges including issues with lab and reagent contamination, allelic dropout, GC content bias, and uneven read distribution which can lead to poor genome recovery (Ospino et al. 2024; Woyke et al. 2011; Lu et al. 2019). Furthermore, MDA generates a large proportion of chimeric artifacts caused by mispriming of displaced strands, which results in artefactual sequences spanning discontinuous regions of the template DNA (Lasken and Stockwell 2007). Chimeric sequencing reads can result in subsequent genome misassembly and false inference of structural variants (Lu et al. 2019). MDA generates two different types of chimeras: direct chimeras and inverted chimeras. Direct chimeras form when a displaced 3’-end re-anneals to a different region of the same template, whereas inverted chimeras form when a displaced 3’-end anneals to a different template strand (Lasken and Stockwell 2007). Previous studies have shown that up to 78% of sequenced MDA products contain chimeras (Lu et al. 2023a; Kiguchi et al. 2021), with inverted chimeras being the most abundant type making up to 91% of detected chimeras (Lu et al. 2023a, 2019). Furthermore, it has been demonstrated that chimeras accumulate during amplification, i.e. greater amplification folds result in higher proportions of chimeric products (Lu et al. 2023a).

Alternative approaches based on MDA have been developed, such as primary template-directed amplification (PTA), which reports more uniform genome coverage (Gonzalez-Pena et al. 2021). However, such approaches yield shorter amplification products, potentially limiting their utility for long-read sequencing. As it stands, MDA offers a reasonable compromise in terms of genome coverage and amplicon fragment size. Recently, MDA has enabled Oxford Nanopore Technologies (ONT) and PacBio long-read sequencing of DNA isolated from individual microscopic animals to generate highly complete genome assemblies (Lee et al. 2023; Roberts et al. 2024). Droplet-based MDA (dMDA) has also been used for PacBio sequencing of DNA isolated from single human cells (Hård et al. 2023). The authors reported that, on average, each read had 2.7 separate alignments which indicates the presence of substantial levels of chimeric artifacts. Although dMDA offered more even genome coverage than conventional MDA, the resulting de novo human genome assembly was less than 20% complete (Hård et al. 2023).

Existing approaches to handle MDA chimeric artifacts are typically based on mapping reads against a reference genome assembly, for example the 3rd-ChimeraMiner tool (Lu et al. 2023a). Such approaches are not suitable for the analysis of novel genomes or de novo genome assembly. Alternatively, tools such as CADECT (Concatemer by Amplification DEteCtion Tool) attempt to identify and remove chimeric reads through read self-alignment (Agyabeng-Dadzie et al. 2025), which risks misclassifying repetitive sequences as chimeras and missing more complex chimeras that cannot be identified from single reads. Several de novo genome assembly algorithms have been developed to assemble short-read MDA sequencing data such as Velvet-SC (Chitsaz et al. 2011), IDBA-UD (Peng et al. 2012), and SPAdes (Nurk et al. 2013a; Bankevich et al. 2012), which include heuristics to mitigate noisy coverage and chimeric reads. To date, there are no long-read genome assembly algorithms designed to handle the extreme amplification bias and high proportions of chimeric reads generated by MDA. Published studies that assemble long-read sequencing of MDA amplified DNA rely on conventional long-read assembly algorithms designed for bulk data. A comprehensive analysis of the impact of chimeric reads on genome assembly structural accuracy is lacking.

Here, we systematically explore the potential and limitations of long-read genome sequencing of MDA-amplified DNA from single cells. We performed long-read sequencing of bulk and single-cell libraries generated from a laboratory culture of *C. reinhardtii*, confirming that MDA-based libraries have extremely high levels of chimeras impacting up to 70% of reads. We show that these chimeric reads lead to the false inference of thousands of structural variants, rendering MDA-derived sequencing data unsuitable for structural variant analysis. Furthermore, de novo assembling these long-read MDA sequencing reads with conventional assembly algorithms generates assemblies with high rates of structural misassembly. To overcome these challenges, we developed lrSAGA (**l**ong-**r**ead **S**ingle **A**mplified **G**enome **A**ssembly), a novel assembly tool to assemble long-read MDA sequencing reads. Assemblies generated using lrSAGA have substantially reduced misassemblies and high base-level accuracy. We apply lrSAGA to long-read genome sequencing of four protist cells isolated from environmental water samples generating high-quality draft genomes. The four cells represent four distinct species: a *Naegleria* amoebaflagellate, a *Bodo* flagellate, and two deep-branching flagellates from the CRuMs supergroup, *Collodictyon triciliatum* and *Diphylleia rotans*. Our work demonstrates the potential of long-read single-cell genomics to generate high-quality draft genomes and explore the genomic diversity of uncultivated microbial eukaryotes.

## Results

### A reference genome assembly for *Chlamydomonas reinhardtii* strain CCAP 11/32A

We aimed to generate a high-quality genome assembly for *C. reinhardtii* strain CCAP 11/32A (equivalent to strains UTEX 90 and SAG 11-32b) to serve as a reference to evaluate sequencing libraries derived from MDA products. Bulk long-read DNA sequencing was performed on a PacBio Revio instrument, yielding approximately 70.7 Gb of HiFi reads with a mean read length of 15 kb, corresponding to an estimated coverage of 617x. HiFi reads were downsampled by discarding reads shorter than 20 kb resulting in an estimated coverage of 90.5x. De novo genome assembly was performed with hifiasm. Following curation, the final nuclear genome assembly comprised 47 contigs totalling 115.2 Mb in length, with a contig N50 of 6.3 Mb and a BUSCO completeness score of 99.7% complete (**Table 1**). We assembled the mitochondrial and plastid genomes separately using Oatk, yielding a linear 15,547 bp mitochondrial genome and a circular 205,825 bp plastid genome. The assembly statistics of our new *C. reinhardtii* CCAP 11/32A assembly compare favourably and has a higher contig N50 than the published reference *C. reinhardtii* CC-4532 v6 genome (Craig et al. 2023) which is 114 Mb in length and has been scaffolded into 17 chromosomes (**Table 1**). The bulk PacBio HiFi reads and genome assembly of *C. reinhardtii* CCAP 11/32A were subsequently used as a reference to assess coverage uniformity and to identify chimeras in the MDA amplified datasets.

**Table 1.** Genome assembly statistics of *Chlamydomonas reinhardtii* CCAP 11/32A and CC-4532 strains.

| Assembly | CC-4532 v6 | CCAP 11/32A |
| --- | --- | --- |
| Reference | Craig et al., 2022 | This study |
| Total length | 114,035,500 bp | 115,168,483 bp |
| Contigs | 120 (17 chromosomes) | 47 |
| Contig N50 | 2,649,501 bp | 6,290,500 bp |
| Scaffold N50 | 6,954,842 bp | - |
| GC (%) | 64.11 | 63.93 |
| BUSCO<br>(Genome mode,<br>Augustus,<br>chlorophyta_odb10) | C:99.6% [S:98.7%,D:0.9%],<br>F:0.1%,<br>M:0.3%,<br>n:1519 | C:99.7%[S:99.2%,D:0.5%],<br>F:0.0%,<br>M:0.3%,<br>n:1519 |
| Mitochondrial genome (linear) | 15,789 bp | 15,547 bp |
| Plastid genome (circular) | 205,535 bp | 205,825 bp |

### Short-read single-cell genome sequencing of *Chlamydomonas reinhardtii*

*C. reinhardtii* CCAP 11/32A cells were sorted into a 96-well plate using fluorescence-activated cell sorting (FACS) (**Fig. S1A**). No cells were sorted into the wells of row A which were used as a negative control. Two *C. reinhardtii* cells were sorted into each well of row B and a single *C. reinhardtii* cell was sorted into each well of rows C to G. A single human cell (K562 cell line) was sorted into each well of row H, for comparison. MDA amplification was performed using the QIAGEN Repli-g Single Cell Kit, followed by short-read Illumina sequencing to generate shallow-coverage sequencing data for quality control.

Excluding a single well that yielded no reads, wells with a single *C. reinhardtii* cell (rows C to G) yielded 0.4 to 3.9 million read pairs (mean 1.4 million) (**Fig. S2A**). The number of read pairs generated from wells with two *C. reinhardtii* cells (row B) ranged from 0.9 million to 2.2 million (mean 1.2 million) (**Fig. S2A**). Wells with a single human cell (row H) generated a comparable number of read pairs, ranging from 1 million to 1.9 million (mean 1.4 million) (**Fig. S2A**). Similarly, wells in row A that contained no cells (negative controls) yielded a similar number of read pairs, ranging from 1 million to 1.6 million (mean 1.4 million) (**Fig. S2A**). Taxonomic classification of reads using Kraken2 classified an average of 45.9% of reads (range 2.8% to 83.5%) from wells with a single *C. reinhardtii* cell as “Chlorophyta” (**Fig. S2B**). For wells with two *C. reinhardtii* cells, the average number of reads classified as “Chlorophyta” was higher at 62.6% (range 45.4% to 80.3%) (**Fig. S2B**). In comparison, for wells containing single human cells, Kraken2 classified on average 98.6% (range 95.6% to 99.1%) of reads as “Human” (**Fig. S2C**). For negative control wells with no sorted cells (i.e. row A), Kraken2 classified on average 95.9% (range 90.2% to 98.1%) of reads as “Human”, indicating contamination. Contaminating “Human” reads were observed in all sequenced wells, with an average of 47.8% (range 12.3% to 93.6%) of reads from wells containing *C. reinhardtii* cells classified as “Human”. The proportion of reads classified as “Human” from wells with two *C. reinhardtii* cells was lower than those from wells with a single *C. reinhardtii* cell (average of 35% versus 50.5%) (**Fig. S2C**). The recovery of human sequences from wells across the entire plate suggests background or reagent contamination.

We estimated the complexity of Illumina libraries by counting unique k-mers and plotting them as a function of the number of reads (**Fig. S3**). Comparing the fraction of unique k-mers at one million reads, wells containing single human control cells showed the highest k-mer diversity (average 77.1% uniqueness), followed by wells containing two *C. reinhardtii* cells (average 62.9% uniqueness) (**Fig. S3**). At this depth, wells containing a single *C. reinhardtii* cell showed similar diversity (average 60.6% uniqueness) (**Fig. S3**). Negative control wells containing no cells showed the lowest diversity (average 51.2% uniqueness) (**Fig. S3**) which was substantially lower than the wells containing human or *Chlamydomonas* cells. The pattern of human contamination across the entire plate, along with the low sequence diversity of contaminated negative controls, and the fact that wells with two *Chlamydomonas* cells have lower levels of contamination (potentially due to having higher amounts of input target DNA) further supports that the contamination source is background or reagent contamination. This highlights the challenges with single-cell genomics. Despite UV treatment to decontaminate materials and performing manual steps within a laminar flow hood, we still observed substantial contamination which consumed a large proportion of sequencing data, reducing the amount of usable data.

Based on the contamination levels (**Fig. S2**) and estimated sequence diversity (**Fig. S3**) from the Illumina skim-sequencing data, six *C. reinhardtii* samples were selected for PacBio HiFi sequencing (below). We selected three single-cell samples (wells F3, E9, and G9) and three double-cell samples (wells B1, B7, and B9) to compare the impact of different starting amounts of template DNA. Additionally, these six samples were subjected to deeper Illumina sequencing to facilitate comparison of short-read and long-read based de novo genome assemblies. The six original Illumina libraries were pooled and resequenced, yielding 4.6 to 8.5 Gb (mean 6 Gb) data, or 15.3 million to 28.4 million read pairs (mean 20.1 million) per sample.

### Long-read single-cell genome sequencing of *Chlamydomonas reinhardtii*

DNA from the six selected samples was divided into two aliquots. One of each of the aliquots was subjected to S1 nuclease digestion. A total of 12 PacBio HiFi libraries (six “MDA” and six “MDA + S1 nuclease”) were pooled and sequenced over two PacBio Revio SMRT cells (**Table S1**). Total HiFi yields for the six undigested MDA libraries ranged from approximately 5 Gb to 5.7 Gb (mean 5.3 Gb) (**Fig. 1A**). S1 nuclease digestion led to a substantial increase in yield with libraries yielding between 7.1 Gb to 10.5 Gb of HiFi data (mean 8.5 Gb) (**Fig. 1A**). This corresponds to a 1.4x to 2.1x increase in yield (mean 1.6x increase) when comparing paired libraries with and without S1 nuclease digestion (**Fig. 1A**). The mean read length of HiFi reads from the undigested MDA libraries ranged from 7.4 kb to 8 kb (**Fig. 1B**). Following S1 nuclease digestion, mean read lengths per library ranged from 6.7 kb to 7.2 kb (**Fig. 1B**), corresponding to a decrease in mean read length by 425 bp to 930 bp per sample. We identified chimeric reads based on mapping against the bulk *C. reinhardtii* CCAP 11/32A genome assembly. For undigested MDA libraries, between 66.6% and 70% reads were identified as being putatively chimeric (**Fig. 1C**). This is in line with previous reports of MDA products having high proportions of chimeras (Lu et al. 2023b). For comparison, upon mapping the bulk HiFi reads back against the assembly, only 0.8% of reads had chimeric alignments. Digestion of MDA products with S1 nuclease reduced the proportion of chimeric reads by 6.2% on average. However, the resulting libraries still exhibited a high proportion of chimeric reads, in the range of 59.8% to 63.4% of reads (**Fig. 1C**). Thus, while S1 nuclease digestion of MDA products only had a minor impact on reducing the proportion of chimeric reads, it proved valuable by substantially increasing overall sequencing yields. The proportion of the *C. reinhardtii* CCAP 11/32A reference genome that was sampled with at least 1x coverage ranged from 68% to 89.2% (**Fig. 1D**). S1 nuclease digested samples exhibited higher completeness than their undigested counterparts; however, downsampling analysis attributed this to their increased sequencing yield (**Table S2**). Double-cell samples generally had higher completeness of the reference genome. However, the overall best sample was a single-cell sample (well G9) (**Fig. 1D**), which had the highest overall completeness of the reference genome. This was also the least contaminated sample (**Fig. S2**). Contamination levels were similar between PacBio data and the corresponding Illumina data (**Fig. S4**), supporting that contamination wasn’t introduced during library preparation. Contaminating human reads were discarded from the following analyses.

**Figure 1.**
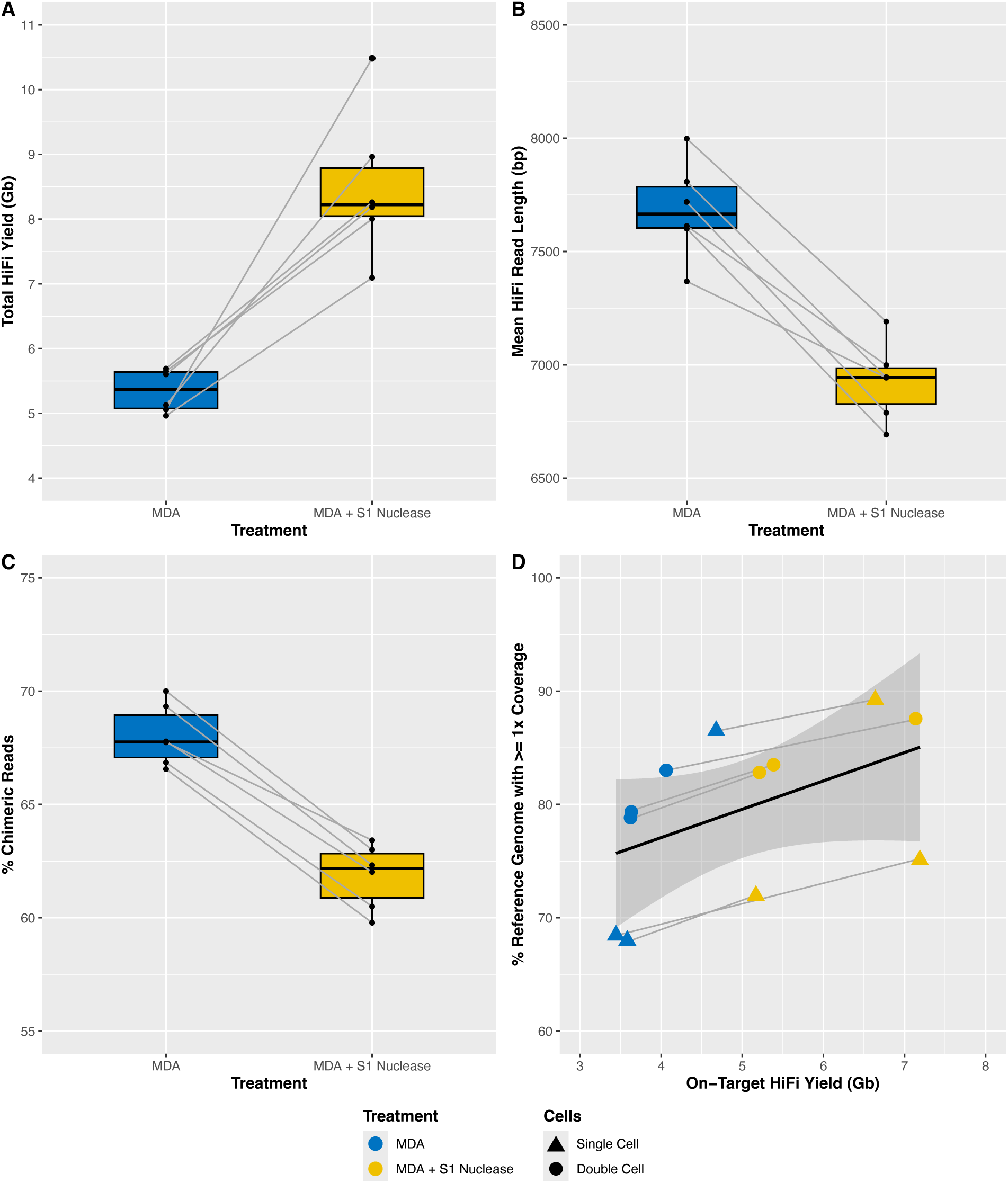
Sequencing statistics of PacBio HiFi libraries generated from MDA amplified DNA from *Chlamydomonas reinhardtii* CCAP 11/32A with and without S1 nuclease digestion. **(A)** Total yield of PacBio HiFi data per sample. **(B)** Mean PacBio HiFi read length per sample. **(C)** Percentage of reads that contain putative chimeras per sample. **(D)** Percentage of the reference genome that was sampled with at least 1x coverage based on mapping reads against the bulk genome assembly, relative to the on-target HiFi yield (i.e. decontaminated) and number of input cells. Grey lines connect libraries that were generated from the same DNA sample with and without S1 nuclease digestion.

Overall, coverage uniformity was very poor across all MDA libraries, regardless of sequencing platform (**Fig. S5A, Fig. S5B**). Note that the high copy number of the plastid genome further skews coverage uniformity estimates (**Fig. S5B**). Although the plastid genome represents only approximately 0.18% of the total reference assembly at 206 kb, it accounts for roughly 16% of the total bulk sequencing reads. Coverage dropouts and regions of extremely high coverage are widespread across the genome (**Fig. S6)**. Regions of extreme high coverage and coverage dropout were observed across all samples (**Fig. S7)**. The poor coverage uniformity is in line with previous single-cell genome sequencing studies that use MDA or MDA-based amplification methods (Macaulay et al. 2015).

Caution is needed when analysing MDA data, as the high proportion of chimeric reads can result in the false inference of structural variants. To explore the extent of this in our data, we called structural variants using Sniffles2 (Smolka et al. 2024) after mapping reads from the PacBio HiFi libraries generated from bulk and MDA amplified DNA against the reference genome assembly of C. reinhardtii CCAP 11/32A. As expected, we did not identify any inversions or duplications in the bulk sequencing data (**Fig. 2**). However, a substantial number of false structural variants were called in each MDA library. Falsely introduced structural variants were dominated by inversions, with between 5,124 and 7,604 inversions called per MDA library (**Fig. 2**). The length of inversions was comparable across all samples, with an overall mean inversion length of 1,493 bp (**Fig. S8**). MDA also introduced false duplications, with 19 to 67 duplications called in each MDA library (**Fig. 2**). Furthermore, MDA elevated the number of insertions and deletions (**Fig. 2**). Overall, S1 digested MDA libraries had a greater number of structural variants compared to untreated MDA libraries, however this was due to the increased yield from S1 digestion. When normalised by read counts, untreated MDA libraries had 1.3 to 1.8-fold more inversions, 2.6 to 4.9-fold more insertions, and 1.3 to 2.2-fold more deletions compared to MDA libraries that were digested with S1 nuclease (**Table S3**).

**Figure 2.**
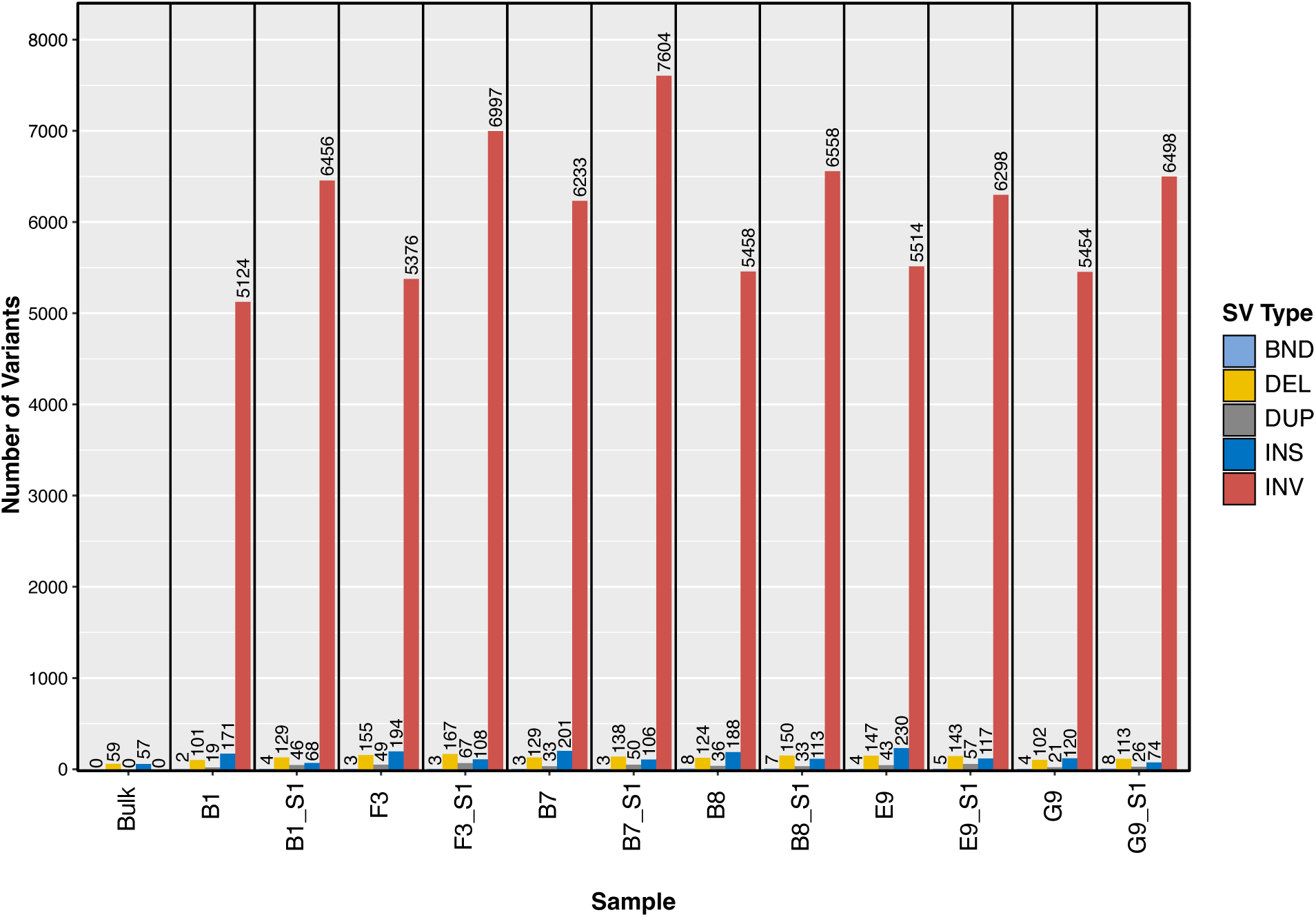
Structural variants identified in *Chlamydomonas reinhardtii* CCAP 11/32A PacBio HiFi libraries generated from bulk or multiple displacement amplified DNA, demonstrating that multiple displacement amplification introduces a substantial number of false structural artifacts. Types of structural variants are indicated: Breakend Translocation (BND), Deletion (DEL), Duplication (DUP), Insertion (INS), and Inversion (INV). See text for corresponding sample IDs.

### De novo genome assembly of MDA amplified DNA from single cells

We speculated that the high proportion of chimeric reads produced by MDA would generate substantial structural errors when assembled using conventional assembly algorithms designed for bulk sequencing data. To test this, we benchmarked the performance of long-read genome assemblers (hifiasm and Flye) and long-read metagenome assemblers (hifiasm-meta, metaFlye, and metaMDBG). Assemblies generated from short-read Illumina sequencing using SPAdes in single-cell mode were included as a baseline for comparison. We used QUAST (Gurevich et al. 2013) to compare each assembly against the reference genome assembly of *C. reinhardtii* CCAP 11/32A. Below, we discuss the results for the best sample (single cell G9/G9_S1). Assembly statistics for all samples are provided in **Fig. 3** and **Table S4**.

**Figure 3.**
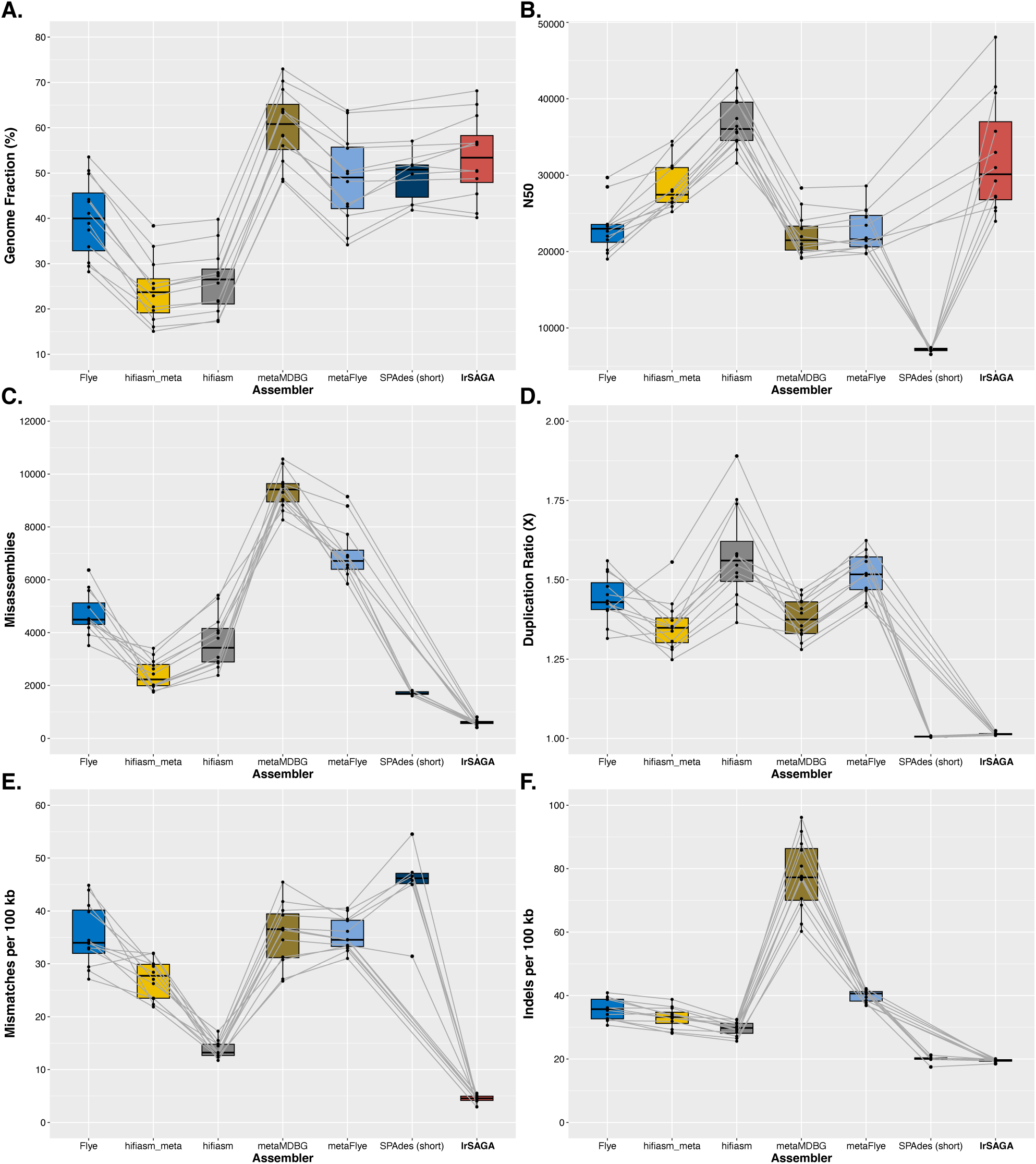
Statistics of *Chlamydomonas reinhardtii* CCAP 11/32A de novo genome assemblies generated from MDA amplified DNA, comparing the performance of long-read assemblers Flye, hifiasm-meta, hifiasm, metaMDBG, metaFlye, lrSAGA, and the short-read assembler SPAdes. Note that there are only six SPAdes assemblies compared to twelve for each long-read assembler (which additionally include S1 nuclease digested DNA). Assembly statistics were computed using QUAST relative to our reference CCAP 11/32A assembly generated from bulk PacBio HiFi sequencing and show genome fraction (**A**), contig N50 (**B**), the number of misassemblies (**C**), duplication ratios (**D**), mismatches per 100 kb (**E**), and indels per 100 kb (**F**).

The SPAdes assembly for sample G9 was 57% complete, with an N50 of 6.5 kb, and contained 1,607 missassemblies and 75 local misassemblies (**Table 2**). For long-read assemblies of sample G9_S1, hifiasm achieved the highest contiguity (N50 = 43.7 kb) but exhibited low completeness (39.8%) and a high number of misassemblies (3,987) (**Table 2**). The metagenome version hifiasm-meta performed similarly but with fewer misassemblies (2,761) (**Table 2**). Flye produced a more complete assembly (53.6%) with an N50 of 29.7 kb but generated even more misassemblies (4,526) (**Table 2**). The metagenome version metaFlye yielded a more complete assembly (63.8%) but also introduced more missassemblies (6,896) and local misassemblies (439) (**Table 2**). metaMDBG generated the most complete assembly (73%) but at the cost of substantially more misassemblies (8,264) and an elevated indel rate (60.24 per 100 kb) (**Table 2**). All long-read assemblies of sample G9_S1 had a high duplication ratio ranging from 1.3x to 1.4x (**Table 2**). Assemblies of other samples showed even higher duplication ratios of up to 1.9x (**Fig. 3**, **Table S4**), indicating that genomic regions were represented multiple times in assemblies which could be falsely interpreted as duplications. The trends were similar across all samples and assemblies (**Fig. 3**). Long-read assemblies had substantially more misassemblies compared to SPAdes short-read assemblies, highly variable levels of completeness, elevated indel rates, and high duplication ratios (**Fig. 3**, **Table S4**).

**Table 2.** De novo genome assembly statistics for *Chlamydomonas reinhardtii* single-cell sample G9_S1.

| Assembler | SPAdes* | hifiasm | Flye | hifiasm-meta | metaFlye | metaMDBG | lrSAGA |
| --- | --- | --- | --- | --- | --- | --- | --- |
| Data Type | Illumina | PacBio HiFi | PacBio HiFi | PacBio HiFi | PacBio HiFi | PacBio HiFi | PacBio HiFi |
| Total length | 66,437,803 | 65,742,721 | 83,436,192 | 56,790,275 | 104,756,371 | 109,687,040 | 79,465,812 |
| Contigs | 16,077 | 1,765 | 4,612 | 1,910 | 5,734 | 5,612 | 3,368 |
| Genome fraction | 57.06% | 39.79% | 53.56% | 38.35% | 63.80% | 72.97% | 68.15% |
| Duplication ratio | 1.003 | 1.422 | 1.344 | 1.28 | 1.415 | 1.28 | 1.009 |
| N50 | 6,549 | 43,733 | 29,693 | 34,434 | 28,593 | 28,328 | 48,076 |
| Largest contig | 61,450 | 215,758 | 190,590 | 174,711 | 329,571 | 222,011 | 361,441 |
| Mismatches per 100 kbp | 31.43 | 12.6 | 27.1 | 23.52 | 32.5 | 26.73 | 4.06 |
| Indels per 100 kbp | 17.5 | 26.5 | 30.62 | 28.44 | 37 | 60.24 | 18.89 |
| Misassemblies | 1,607 | 3,987 | 4,526 | 2,761 | 6,896 | 8,264 | 415 |
| Local misassemblies | 75 | 90 | 311 | 130 | 439 | 390 | 15 |
| Consensus Quality (QV) | 36 | 34.1 | 35.4 | 36.8 | 34.4 | 30.3 | 41 |

370 \*The SPAdes assembly shown is from sample G9 (i.e. not treated with S1 nuclease).

We also benchmarked CADECT, a tool to identify and remove concatemeric sequences. CADECT segments reads into windows and aligns different windows from the same read against each other to detect chimeras (Agyabeng-Dadzie et al. 2025). For sample G9_S1, CADECT classified only 26.2% of reads as “putative concatemers”. This is substantially less than our estimate of 60.5% based on mapping against the reference genome (**Fig. 1, Table S1**). Upon mapping reads identified by CADECT as “non-concatemers” against our reference genome assembly, 56.8% of the reads were classified as chimeric based on our criteria above.

Furthermore, assembling the reads classified by CADECT as “non-concatemers” resulted in only a modest reduction in the number of misassemblies (16.4% fewer misassemblies on average), but this came at the cost of smaller assemblies (18.31% smaller on average) with lower genome completeness (16.75% lower completeness on average) with similar duplication ratios (**Table S4)**. These results demonstrate that existing assembly algorithms designed for bulk sequencing data perform poorly on libraries derived from single-cell MDA amplified DNA, and that current tools for handling chimeric reads are inadequate.

We developed lrSAGA, a tool for **l**ong-**r**ead **S**ingle **A**mplified **G**enome **A**ssembly designed to address the high chimera rate and noisy coverage characteristic of MDA amplified DNA. lrSAGA takes as input a raw, unprocessed assembly graph in GFA format. Here, we use MBG (Rautiainen and Marschall 2021) to construct the initial assembly graph. MBG performs homopolymer compression of input reads, generates a de Bruijn Graph using syncmers, and outputs the graph in GFA format with homopolymer runs expanded. Like most assembly approaches, lrSAGA progressively simplifies the assembly graph to generate contigs, iteratively removing tips, bubbles, transitive edges, and chimeric sequences, and merges unambiguous linear paths to form contigs. Chimeric segments can present with relatively high sequencing depth if the chimeric fragment forms early during the amplification process and subsequently serves as a template for further amplification. Thus, discarding low coverage nodes is insufficient for detecting chimeric sequences. Furthermore, simply discarding low coverage nodes risks discarding bona fide genomic sequences due to the extreme coverage heterogeneity introduced by MDA (Nurk et al. 2013b). lrSAGA identifies chimeras based on assembly graph topology. Chimeric nodes interrupt otherwise linear paths through the assembly graph. Inverted chimeras are identifiable as nodes that overlap two neighbouring nodes, one in the forward orientation and the other in the reverse complement orientation (**Fig. S9**). For example, given three nodes X, Y, and Z that overlap in the same orientation giving X→Y→Z but are interrupted by a chimeric node C, whereby C partially overlaps with X in the forward orientation but the reverse complement of C also partially overlaps with Y. In lrSAGA, putative inverted chimeras are identified as nodes with two neighbours that also overlap with each other (**Fig. S9**). Provided sufficient non-chimeric reads span a particular genomic region, it should be possible to assemble the correct sequence regardless of the depth of chimeric reads. lrSAGA is implemented in Python and makes use of several external libraries including GfaPy (Gonnella and Kurtz 2017) for GFA file processing and NetworkX (Hagberg et al. 2008) for graph traversal. lrSAGA parameters are exposed to the user, for example, to conservatively limit node removal to short low-coverage nodes. Our approach can be applied to raw PacBio HiFi reads, however, better performance can be expected when run on error-corrected HiFi reads. Here, we used hifiasm to correct errors in PacBio HiFi reads before constructing an assembly graph with MBG. lrSAGA can also be applied to high-quality or error-corrected ONT reads (see below).

We benchmarked lrSAGA using the *C. reinhardtii* MDA sequencing reads. The lrSAGA assembly of sample G9_S1 outperformed other assemblies across most metrics. It had the highest contig N50 at 48 kb despite our conservative assembly approach and, unlike other long-read assemblies, maintained a low duplication ratio (1.009x) (**Table 2**). It was also the most accurate at the base level overall in terms of mismatches (4.06 per 100 kb) and its indel rate (18.9 per 100 kb) was comparable to that of the Illumina assembly (**Table 2**). We also estimated consensus quality values (QV) separately using Merqury, which was estimated to be Q41 (99.992% accuracy) for the lrSAGA assembly (**Table 2**). Importantly, lrSAGA generated substantially fewer misassemblies than all other approaches (**Table 2**). With only 415 misassemblies and 15 local misassemblies, the lrSAGA assembly had 85% fewer misassemblies relative to the next best long-read assembly and 74% fewer misassemblies than even the SPAdes short-read assembly. Despite the reduction in structural errors, lrSAGA maintained a high genome completeness of 68.15% (**Table 2**). This completeness is close to that of metaMDBG but with 95% fewer misassemblies.

These trends were similar across all 12 *C. reinhardtii* samples. Assemblies generated by lrSAGA were amongst the most complete (**Fig. 3A)** and most contiguous (**Fig. 3B**) compared to other assembly algorithms. Most significantly, lrSAGA assemblies had substantially reduced misassemblies compared to all other assemblers, including SPAdes (**Fig. 3C**). Other long-read assemblers generated assemblies with inflated assembly lengths (**Table S4**) relative to genome completeness corresponding to high duplication ratios (**Fig. 3D**). This was not the case for lrSAGA. lrSAGA assemblies were also the most accurate overall at the base-level in terms of mismatches (**Fig. 3E**). Likewise, lrSAGA outperformed other long-read assemblers with substantially reduced rates of indels, comparable to that of SPAdes assemblies of Illumina data (**Fig. 3F**). lrSAGA assemblies also had the highest QV scores overall, with an average of Q40 (range Q38.5 – Q41.2) (**Table S4**) corresponding to an average base accuracy of 99.99%. Taking all metrics into account, lrSAGA achieved the best balance overall in terms of completeness and contiguity while maintaining high structural and base-level accuracy. While the widespread coverage dropouts (**Fig. S6, Fig. S7**) limits contiguity for all assemblers, it is remarkable that we can generate an accurate de novo genome assembly with up to 68% completeness from just a single haploid cell as input.

### De novo genome assembly of MDA amplified DNA from microscopic animals

To further validate our assembly approach, we reanalysed published MDA sequencing reads from the nematode *Caenorhabditis elegans*. Roberts et al. (2024) performed PacBio HiFi sequencing of MDA amplified DNA isolated from one-half of an individual *C. elegans* worm (Bristol N2 strain). We downloaded the PacBio HiFi reads (SRR27483171) and generated a genome assembly using hifiasm for comparison. Hifiasm generated a 121.4 Mb assembly across 493 contigs with a contig N50 of 559 kb (**Table 3**). We compared this assembly to the reference *C. elegans* Bristol N2 genome GCF_000002985.6 (The *C. elegans* Sequencing Consortium 1998) using QUAST. The hifiasm assembly was 96.85% complete but larger than expected with a high duplication ratio of 1.24 (**Table 3**). QUAST identified a large number of structural errors with 1,753 misassemblies and 487 local misassemblies (**Table 3**). In terms of base-level accuracy, 21.4 mismatches per 100 kb and 17.4 indels per 100 kb were identified (**Table 3**). To benchmark our assembly process, we generated a de novo assembly using lrSAGA. The lrSAGA assembly was 100.1 Mb in length across 353 contigs, with a higher contig N50 of 795 kb (+42.2%) (**Table 3**). Genome completeness was slightly higher at 98.6% (+1.7%) (**Table 3**). The lrSAGA assembly did not suffer from false duplications with a duplication ratio of 1.01x (**Table 3**). Importantly, the lrSAGA assembly was substantially more accurate at the structural-level with 329 misassemblies (−81.2%), 423 local misassemblies (−13%), and also at the base-level with 19.7 mismatches per 100 kb (−7.8%), and 7.9 indels per 100 kb (−54.4%) (**Table 3**).

**Table 3.**
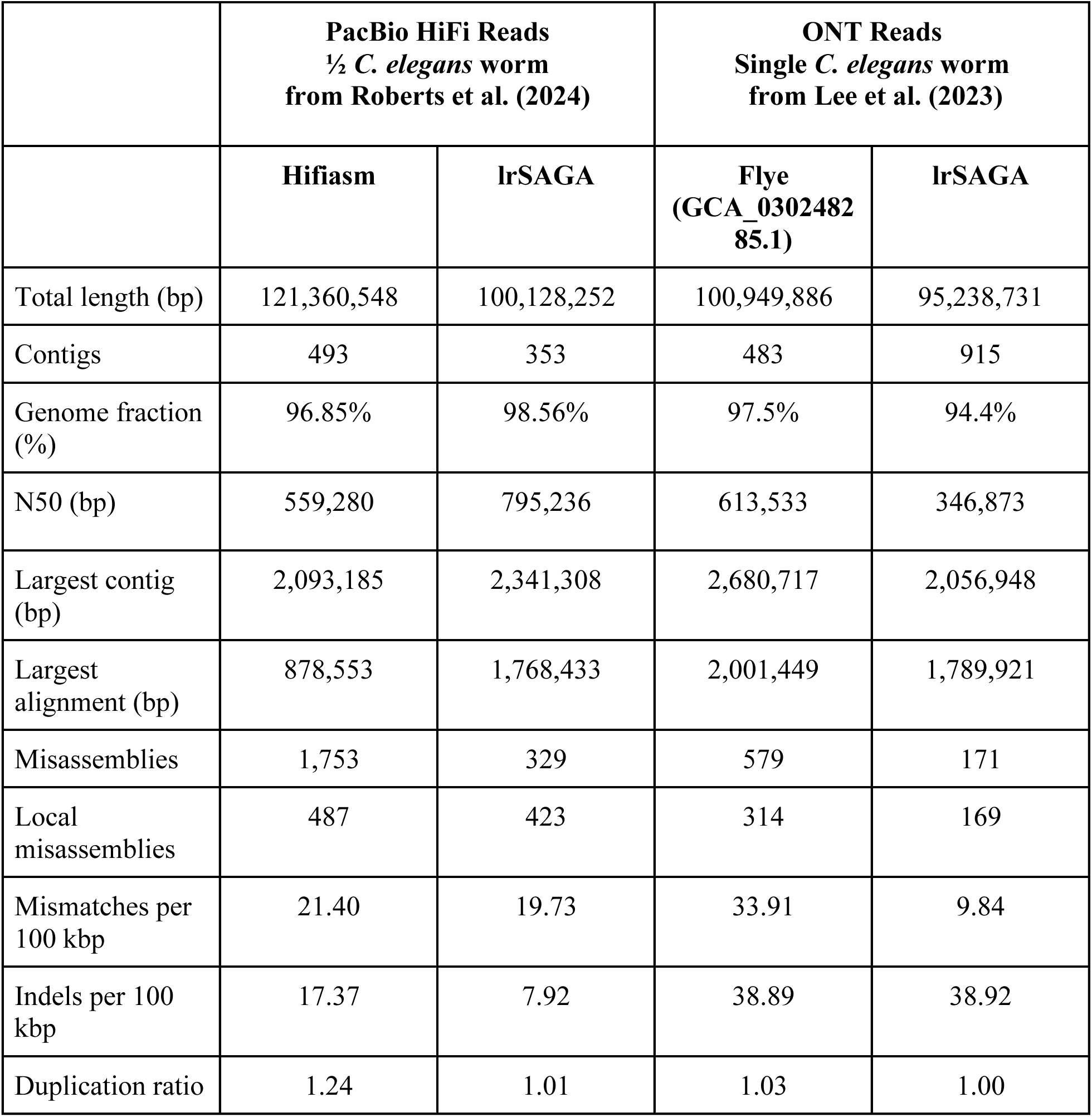
De novo genome assembly statistics for *Caenorhabditis elegans* MDA amplified DNA.

|  | **PacBio HiFi Reads**<br>**½ *C. elegans* worm**<br>**from Roberts et al. (2024)** |  | **ONT Reads**<br>**Single *C. elegans* worm**<br>**from Lee et al. (2023)** |  |
| --- | --- | --- | --- | --- |
|  | **Hifiasm** | **lrSAGA** | **Flye**<br>**(GCA_0302482**<br>**85.1)** | **lrSAGA** |
| Total length (bp) | 121,360,548 | 100,128,252 | 100,949,886 | 95,238,731 |
| Contigs | 493 | 353 | 483 | 915 |
| Genome fraction (%) | 96.85% | 98.56% | 97.5% | 94.4% |
| N50 (bp) | 559,280 | 795,236 | 613,533 | 346,873 |
| Largest contig (bp) | 2,093,185 | 2,341,308 | 2,680,717 | 2,056,948 |
| Largest alignment (bp) | 878,553 | 1,768,433 | 2,001,449 | 1,789,921 |
| Misassemblies | 1,753 | 329 | 579 | 171 |
| Local misassemblies | 487 | 423 | 314 | 169 |
| Mismatches per 100 kbp | 21.40 | 19.73 | 33.91 | 9.84 |
| Indels per 100 kbp | 17.37 | 7.92 | 38.89 | 38.92 |
| Duplication ratio | 1.24 | 1.01 | 1.03 | 1.00 |

We also validated our assembly approach using ONT sequencing reads. Lee et al. (2023) performed ONT sequencing of MDA amplified DNA isolated from a single *C. elegans* worm (Bristol N2 strain) that was digested with T7 endonuclease. The authors attempted to handle chimeras by splitting reads that map to the reverse complement of themselves. They generated an assembly using Flye and polished the assembly with the ONT reads and also Illumina reads. The resulting assembly was 100.9 Mb across 483 contigs with a contig N50 of 613.5 kb (**Table 3**). We compared the published assembly from Lee et al. (GCA_030248285.1) to the reference *C. elegans* assembly using QUAST. QUAST determined that the assembly was 97.5% complete, and identified 579 misassemblies, 314 local misassemblies, 33.91 mismatches per 100 kb, and 38.89 indels per 100 kb (**Table 3**). We downloaded the ONT reads generated by Lee et al. (SRR24201716). These sequencing reads were generated using older ONT chemistry (R9.4.1) with lower base accuracy (median read quality estimate of Q14 - 96% accuracy) so we performed read error correction using NECAT (Chen et al. 2021), prior to genome assembly. lrSAGA generated a 95 Mb assembly across 915 contigs (**Table 3**). Our assembly was less contiguous than the published assembly with a contig N50 of 347 kb and also less complete at 94.4% complete (**Table 3**). However, lrSAGA did generate a more accurate assembly with only 171 misassemblies (−70.5%), 169 local misassemblies (−46.2%), and only 9.84 mismatches per 100 kb (−71%) (**Table 3**). The indel rate was comparable at 38.92 indels per 100 kb (**Table 3**). We note that the lrSAGA assemblies weren’t polished after assembly. This demonstrates that our approach can also generate accurate de novo genome assemblies from ONT sequencing of MDA amplified DNA.

### Long-read single-cell genome sequencing of environmental protists

To demonstrate our approach with environmental samples, we sequenced four single protist cells isolated from river water. The four cells were recovered via FACS of a sample of surface water collected from the River Leam in Royal Leamington Spa (UK) in August 2022. DNA was isolated from each cell and WGA was performed using MDA as part of a modified G&T-seq protocol (Macaulay et al. 2015). Note that these DNA samples were not digested with S1 nuclease. The four libraries were pooled and sequenced on one PacBio Sequel IIe SMRT Cell.

PacBio HiFi reads were error corrected using hifiasm and then assembled with MBG and lrSAGA. If present, sequences from bacterial co-bionts were manually removed based on a combination of binning with MetaBat2 and taxonomic classification using CAT and Tiara. The identity of the cells were determined post-sequencing based on analysis of the small subunit (SSU) ribosomal RNA genes recovered from the assembled genomes. The four cells were identified as *Collodictyon triciliatum*, *Diphylleia rotans*, and unnamed members of the *Bodo* and *Naegleria* genera.

We previously sequenced the same *Naegleria* DNA sample (referred to as *Naegleria* sp. PL0398), identified as a close relative of *N. pagei*, and generated an Illumina-based assembly (McGowan et al. 2025). Here, PacBio HiFi sequencing yielded 4.6 Gb of data across 633,239 reads with a read length N50 of 7.7 kb (**Table 4**). The final lrSAGA assembly, after removing the genome of a Legionellaceae co-biont, was 44.42 Mb in length across 310 contigs with a contig N50 of 253.8 kb (**Table 4**). Average coverage depth, calculated by aligning the PacBio HiFi reads against the final assembly, was determined to be 95x and 88.76% of the assembly was sampled with at least 20X read coverage (**Table 4**). Approximately 8.1% (3.6 Mb) of the assembly was annotated as repetitive (**Table 4**). Genome annotation identified 22,261 protein-coding genes with a BUSCO completeness of 84.3% (**Table 4**). Compared to our previous short-read assembly of this DNA sample, the lrSAGA long-read assembly is approximately 3 Mb larger with a 15-fold improvement in contig N50. The karyotype of this uncultivated *Naegleria* species is not known, however a chromosomal-level genome assembly of *Naegleria fowleri* has been published with 37 chromosomes ranging in length from 537 kb to 1.2 Mb (N50 of 756,811 bp) (Ali et al. 2021). The largest contig in our lrSAGA assembly of *Naegleria* sp. PL0398 is 962.6 kb (**Table 4**) and 10 contigs are larger than 490 kb, supporting that our assembly is highly contiguous. Furthermore, our single-cell assembly is more contiguous than some *Naegleria* genomes that were sequenced from bulk cultures, including long-read based assemblies, such as the ONT based assembly of *Naegleria clarki* (Willemsen et al. 2025).

**Table 4.**
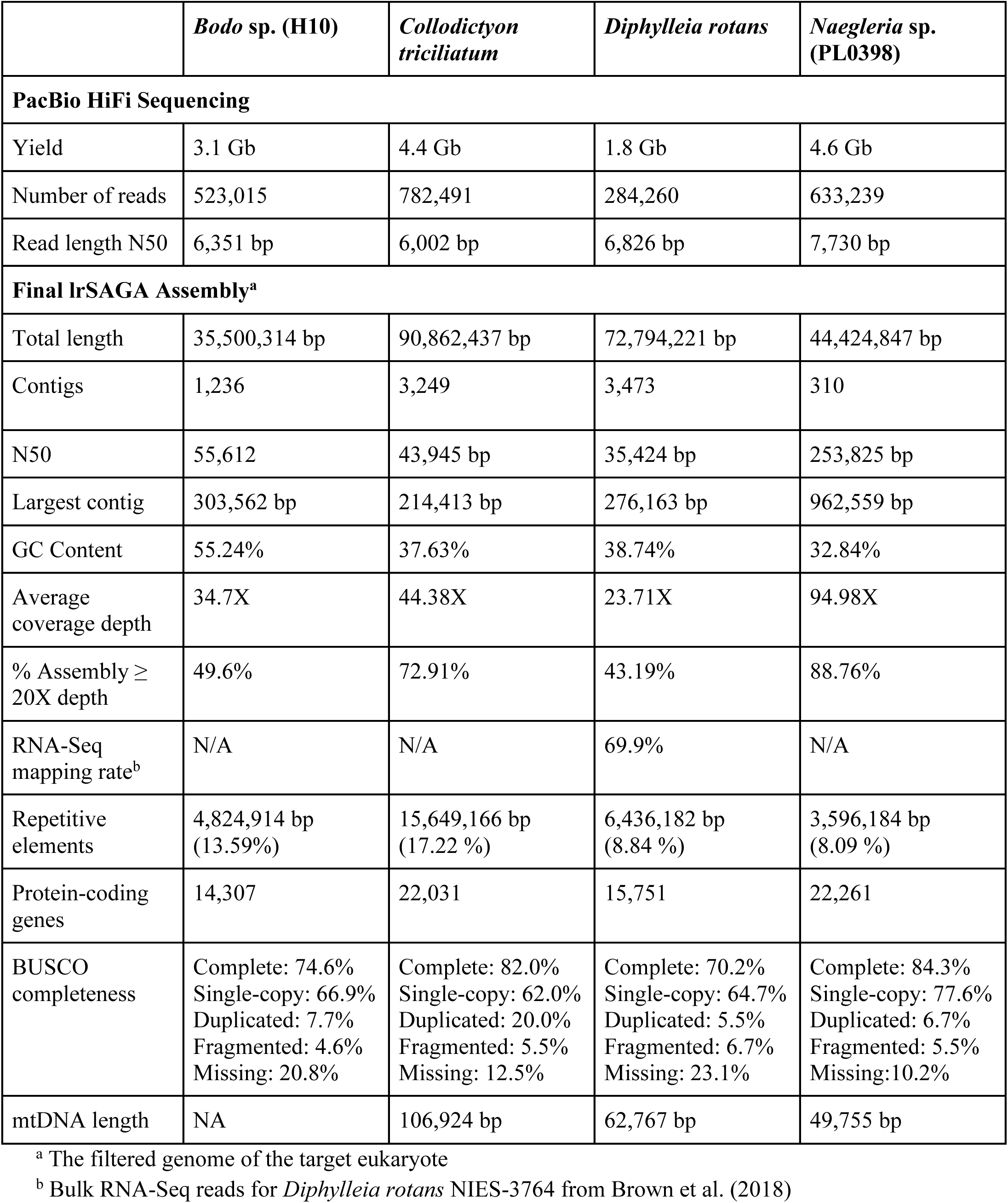
lrSAGA long-read single-cell genome assemblies of four environmental protists.

|  | *Bodo* sp. (H10) | *Collodictyon triciliatum* | *Diphylleia rotans* | *Naegleria* sp. (PL0398) |
| --- | --- | --- | --- | --- |
| **PacBio HiFi Sequencing** |  |  |  |  |
| Yield | 3.1 Gb | 4.4 Gb | 1.8 Gb | 4.6 Gb |
| Number of reads | 523,015 | 782,491 | 284,260 | 633,239 |
| Read length N50 | 6,351 bp | 6,002 bp | 6,826 bp | 7,730 bp |
| **Final lrSAGA Assembly<sup>a</sup>** |  |  |  |  |
| Total length | 35,500,314 bp | 90,862,437 bp | 72,794,221 bp | 44,424,847 bp |
| Contigs | 1,236 | 3,249 | 3,473 | 310 |
| N50 | 55,612 | 43,945 bp | 35,424 bp | 253,825 bp |
| Largest contig | 303,562 bp | 214,413 bp | 276,163 bp | 962,559 bp |
| GC Content | 55.24% | 37.63% | 38.74% | 32.84% |
| Average coverage depth | 34.7X | 44.38X | 23.71X | 94.98X |
| % Assembly ≥ 20X depth | 49.6% | 72.91% | 43.19% | 88.76% |
| RNA-Seq mapping rate<sup>b</sup> | N/A | N/A | 69.9% | N/A |
| Repetitive elements | 4,824,914 bp (13.59%) | 15,649,166 bp (17.22 %) | 6,436,182 bp (8.84 %) | 3,596,184 bp (8.09 %) |
| Protein-coding genes | 14,307 | 22,031 | 15,751 | 22,261 |
| BUSCO completeness | Complete: 74.6%<br>Single-copy: 66.9%<br>Duplicated: 7.7%<br>Fragmented: 4.6%<br>Missing: 20.8% | Complete: 82.0%<br>Single-copy: 62.0%<br>Duplicated: 20.0%<br>Fragmented: 5.5%<br>Missing: 12.5% | Complete: 70.2%<br>Single-copy: 64.7%<br>Duplicated: 5.5%<br>Fragmented: 6.7%<br>Missing: 23.1% | Complete: 84.3%<br>Single-copy: 77.6%<br>Duplicated: 6.7%<br>Fragmented: 5.5%<br>Missing: 10.2% |
| mtDNA length | NA | 106,924 bp | 62,767 bp | 49,755 bp |

<sup>a</sup> The filtered genome of the target eukaryote
<sup>b</sup> Bulk RNA-Seq reads for *Diphylleia rotans* NIES-3764 from Brown et al. (2018)

We also previously sequenced the same *Bodo* DNA sample (*Bodo* H10) and generated an Illumina-based assembly, identifying the cell as a member of the free-living flagellate genus *Bodo* and harbouring a bacterial co-biont (Warring et al. 2026). Here, PacBio HiFi sequencing yielded 3.1 Gb of data across 523,015 reads with a read length N50 of 6.4 kb (**Table 4**). Following removal of the bacterial co-biont genome, the final lrSAGA assembly was 38 Mb in length across 1,378 contigs with a contig N50 of 54.5 kb (**Table 4**). This represents a 2.5-fold improvement in contig N50 compared to our previous Illumina assembly of the same DNA sample (Warring et al. 2026). The average coverage depth was 34.7X; however, only 49.6% of the assembly was sampled with at least 20X read coverage (**Table 4**). This suggests that the low coverage depth may have limited contiguity. Approximately 13.6% of the assembly was annotated as repetitive and 14,307 protein coding genes were predicted with a BUSCO completeness of 74.6% (**Table 4**).

For the cell identified as *Collodictyon triciliatum*, PacBio HiFi sequencing yielded 4.4 Gb of data across 782,491 reads with a shorter read length N50 of 6 kb (**Table 4**). The final lrSAGA assembly, after removing bacterial sequences, was 90.9 Mb in length across 3,249 contigs with a contig N50 of 43.9 kb (**Table 4**). The raw assembly graph for this sample was relatively sparse which is indicative of coverage dropouts and/or low sequencing depth. The average coverage depth was 44x with 72.91% of the assembly sampled with a least 20X read coverage (**Table 4**). A single eukaryotic SSU rRNA gene was identified in this assembly which was 99.67% identical to that of *C. triciliatum* isolate Arungen (MF039367.1). *C. triciliatum* is a cryptic species complex of quadriflagellates that are morphologically indistinguishable under light microscopy (Orr et al. 2018; Galindo et al. 2024). Phylogenetic analysis placed the SSU rRNA sequence from our single cell as sister to *C. triciliatum* isolate Arungen with 90% bootstrap support within the “Diphy II” clade (**Fig. S10**). Approximately 17.2% (15.6 Mb) of the assembly was annotated as repetitive. Genome annotation identified 22,031 protein-coding genes with a BUSCO completeness of 82.0% (**Table 4**).

A lower PacBio HiFi yield was recovered for the cell identified as *Diphylleia rotans*, with 1.8 Gb of data across 284,260 reads with a read length N50 of 6.8 kb (**Table 4**). The final lrSAGA assembly for this sample was 72.8 Mb across 3,473 contigs with a contig N50 of 35.4 kb. Similarly, the raw assembly graph for this sample was less densely connected and the mean average coverage depth of the final assembly was lower at 23.7x (**Table 4**). Only 43.19% of the assembly was sampled with at least 20X read coverage (**Table 4**). A single eukaryotic SSU rRNA gene was identified in this assembly which was 99.7% identical to *D. rotans* isolate NIES3764. Phylogenetic analysis placed the SSU rRNA sequence from our single cell within the “Diphy I” clade of biflagellates classified as *D. rotans* (**Fig. S10**). Furthermore, the mitochondrial genome recovered from the assembly was 99.99% identical and had a full length alignment to the mitochondrial genome of *D. rotans* isolate NIES-3764 (Kamikawa et al. 2016). A bulk RNA-seq dataset was previously published for *D. rotans* isolate NIES-3764 (Brown et al. 2018). Aligning the RNA-seq reads against our single-cell genome assembly resulted in an overall alignment rate of 69.9% (**Table 4**), supporting that our assembly is highly complete. Only 1.95% of RNA-seq read pairs aligned discordantly (i.e., read-pairs that aligned in the incorrect orientation, to different contigs, or with an unexpected insert size) suggesting that our assembly is structurally accurate. Approximately 8.8% (6.4 Mb) of the assembly was annotated as repetitive (**Table 4**). Genome annotation, supported by the bulk RNA-seq data, identified 15,751 protein-coding genes with a BUSCO completeness of 70.2% (**Table 4**).

## Discussion

Multiple displacement amplification has been transformative for single-cell genomics, enabling whole-genome amplification and sequencing of quantities of DNA that are otherwise inaccessible. Its low error rate, high yield, and relatively long amplification products have made it a popular method for single-cell genome sequencing (Spits et al. 2006; Dean et al. 2002). However, MDA is fundamentally limited by several problems, including reagent contamination, uneven amplification, coverage dropouts, and the generation of chimeric products that result in artefactual sequencing spanning discontinuous regions of template DNA (Lu et al. 2023b; Woyke et al. 2011; Ospino et al. 2024). In particular, the high proportion of chimeric products has limited the application of long-read single-cell genome sequencing, with previous studies reporting that up to 78% of long reads derived from MDA products are chimeric (Kiguchi et al. 2021; Lu et al. 2023a). A comprehensive analysis on the impact of chimeric reads on structural variant identification and de novo genome assembly has been lacking. Here, we systematically evaluated these problems in the context of long-read single-cell genome sequencing. Using the model single-celled green *Chlamydomonas reinhardtii*, we benchmarked the performance of PacBio HiFi sequencing of MDA-amplified from single cells, quantifying the consequences of chimeric reads for structural variant calling and de novo genome assembly. We demonstrated that conventional tools designed for bulk sequencing data are not suitable for the analysis of long-read MDA data. We developed lrSAGA, a novel assembly tool to address these issues.

We evaluated the impact of digesting MDA products with S1 nuclease, which cleaves single stranded nucleic acids. We found that digestion of MDA products with S1 nuclease prior to library preparation substantially increased PacBio HiFi sequencing yields. S1 nuclease digested samples yielded on average approximately 1.6x more data (range 1.4x to 2.1x) compared to undigested samples (**Fig. 1A**). This increase in sequencing yield is likely due to the reduction of hyperbranched DNA structures which are characteristic of MDA(Agyabeng-Dadzie et al. 2025; Lu et al. 2023b; Lasken and Stockwell 2007). The increase in sequencing yield led to a higher genome fraction recovery (**Table S2**). This increase in yield was accompanied by a modest reduction in mean read length of 425 to 930 bp per sample compared to undigested samples (**Fig. 1B**). Despite its benefits for sequencing yield, S1 nuclease digestion resulted in only a minor reduction in the proportion of chimeric reads, with digested samples yielding 59.8% to 63.4% chimeric reads compared to 66.6% to 70% for undigested MDA products (**Fig. 1C**). Further reductions in chimera frequency may be achievable through optimisation of S1 nuclease treatment conditions or by combining S1 digestion with additional post-amplification processing steps (Zhang et al. 2006). Thus, while S1 nuclease digestion proved valuable by substantially increasing overall sequencing yields, it does not solve the chimera problem. This problem is evident when MDA amplified DNA is used to call structural variants. Calling structural variants from PacBio HiFi reads derived from MDA-amplified DNA resulted in the false inference of thousands of structural variants per sample compared to bulk sequencing of the same *C. reinhardtii* laboratory culture (**Fig. 2**). Our results demonstrate that structural variant calls derived from MDA-amplified sequencing reads are not reliable and that methods to accurately distinguish structural variants from MDA artefacts are lacking. This has serious implications for clinical and biomedical applications of single-cell MDA, such as studies that attempt to link somatic structural variants to disease phenotypes.

We benchmarked five long-read assembly algorithms (hifiasm, hifiasm-meta, Flye, metaFlye, and metaMDBG) using our *C. reinhardtii* MDA sequencing datasets and found that all performed poorly, producing assemblies with high misassembly rates, inflated duplication ratios, and highly variable genome completeness (**Figure 3**, **Table 2**). Notably, the number of misassemblies in long-read assemblies were substantially elevated compared to short-read assemblies generated by SPAdes from Illumina sequencing of the same DNA samples (**Figure 3, Table S4**). Metagenome assemblers designed to tolerate more variable levels of coverage, in particular metaFlye and metaMDBG, produced more complete assemblies but at the cost of even higher misassembly rates (**Figure 3**, **Table 2**). We note that the long-read assemblers benchmarked here are not designed to handle such high rates of chimeric reads so their poor performance here is not a reflection of their intended purpose. CADECT, which attempts to identify and remove chimeric sequences through read self-alignment, substantially underestimated the true chimera rate and only modestly reduced subsequent misassemblies (**Table S1**). These results demonstrate that assembly algorithms designed for bulk sequencing data are unsuited for MDA datasets and that existing tools to pre-filter chimeric reads are inadequate given the scale of chimeric content in single-cell MDA datasets.

To address these challenges, we developed lrSAGA, a new assembly tool designed for long-read datasets with high chimera rates and noisy coverage. Identifying chimeras based on sequencing coverage alone is insufficient as chimeric sequences can have relatively high sequencing depth if they form early during amplification and subsequently serve as templates for further amplification. Furthermore, simply discarding low-coverage sequences risks losing genuine genomic content due to the extreme coverage heterogeneity introduced by MDA (Bankevich et al. 2012). Instead, lrSAGA identifies putative chimeras based on assembly graph topology. Across all twelve *C. reinhardtii* PacBio HiFi MDA libraries benchmarked, lrSAGA consistently produced assemblies that were among the most complete and contiguous while achieving drastically reduced misassembly rates (**Table 2**, **Figure 3**). For the best single-cell sample (G9_S1), the lrSAGA assembly had 85% fewer misassemblies than hifiasm-meta which was the next best in terms of misassemblies, while maintaining a high genome completeness of 68% (**Table 2**). Compared to metaMDBG, which had the highest completeness of 72.97%, lrSAGA generated 95% fewer misassemblies (**Table 2**). lrSAGA misassemblies were also substantially reduced compared to Illumina-based assemblies generated by SPAdes, a short-read assembler with a single-cell mode designed for MDA datasets (**Figure 3**). lrSAGA assemblies were also the most accurate overall at the base level in terms of mismatches, and indel rates were comparable to SPAdes assemblies of Illumina data (**Figure 3**). Despite the conservative assembly approach, lrSAGA assemblies were also among the most contiguous (**Figure 3**). However, contiguity is ultimately limited by the widespread coverage dropouts (**Figure S6, Figure S7**). Our results are particularly remarkable given that the input data has chimera rates of up to 70%, extreme coverage heterogeneity, and that the input DNA came from just a haploid single cell. Thus, we demonstrated that we can generate highly complete and highly accurate (both at the base and structural level) assemblies with lrSAGA from single cells.

To confirm that lrSAGA generalises beyond our *C. reinhardtii* benchmark dataset, we reanalysed published long-read sequencing data from MDA-amplified DNA derived from single or half of individual *Caenorhabditis elegans* worms. Reassembly of PacBio HiFi reads from Roberts et al. (2024) using lrSAGA yielded a more complete (98.56% complete), more contiguous (795 kb contig N50), and substantially more accurate assembly compared to hifiasm, with 81% fewer misassemblies (**Table 3**). The dataset from Roberts et al. (2024) was generated from just one-half specimen of *C. elegans*. This approach is very promising as one-half of an individual isolated from the environment could be subjected to MDA for genome sequencing, while the other half could be used to generate complimentary data-types, e.g., RNA-sequencing to support genome annotation, low-input Hi-C to scaffold the assembly, or proteomic and metabolomic profiling, with ongoing technological advances continuing to lower sample input requirements. We additionally benchmarked lrSAGA using ONT sequencing reads from Lee et al. (2023), which were generated by MDA of a single *C. elegans* worm. Here, lrSAGA generated a less contiguous (347 kb contig N50) and less complete (94.4% complete) assembly compared to the published assembly (**Table 3**). The poorer performance of lrSAGA is expected given lower quality reads from older ONT chemistry (R9.4.1). We anticipate better performance from more recent ONT chemistries with higher read accuracies. Nevertheless, lrSAGA generated a more accurate assembly with 70.5% fewer misassemblies and 46% fewer local misassemblies (**Table 3**). Furthermore, the lrSAGA was more accurate at the base-level with 71% fewer mismatches (**Table 3**), despite not being polished post-assembly. The *C. elegans* benchmarking demonstrates that lrSAGA is robust across different sequencing platforms and input data qualities.

A key application of MDA is the sequencing of uncultivated microorganisms from environmental samples. Here, we performed long-read genome sequencing of four single protist cells isolated from environmental water sampling. The four cells represent four distinct species: a *Naegleria* amoebaflagellate, a *Bodo* flagellate, and two deep-branching flagellates from the CRuMs supergroup, *Collodictyon triciliatum* and *Diphylleia rotans*. PacBio HiFi libraries were generated from MDA amplified DNA from each single cell, and assembled using lrSAGA, yielding high-quality draft genome assemblies estimated to be 70 - 84% complete by BUSCO (**Table 4**). The *Naegleria* sample performed best overall, yielding an assembly estimated to be 84.3% complete with a contig N50 of 253.8 kb (**Table 4**). This represents a 15-fold improvement in contig N50 compared to a short-read assembly that we previously generated from the same *Naegleria* DNA sample (McGowan et al. 2025). Strikingly, this single-cell assembly is more contiguous than published *Naegleria* genomes that were sequenced from bulk cultures, including a long-read ONT-based assembly of *N. clarki* (Willemsen et al. 2025). The *Bodo* assembly was estimated to be 74.6% complete by BUSCO and had a contig N50 of 55.6 kb (**Table 4**), which is a 2.5-fold improvement over our previous short-read assembly (Warring et al. 2026). For *C. triciliatum* and *D. rotans*, our assemblies represent, to our knowledge, the first genome sequences available for either species. These assemblies were less contiguous with contig N50s of 43.9 kb and 35.4 kb, respectively (**Table 4**), which is likely attributable to lower PacBio HiFi yields relative to genome size resulting in sparser assembly graphs. Despite this, both assemblies were estimated to be highly complete, with BUSCO completeness scores of 82% and 70.2% respectively (**Table 4**). For *D. rotans*, alignment of published bulk RNA-seq reads (Brown et al. 2018) against our single-cell assembly yielded an overall alignment rate of 69.9% with only 1.95% discordant read pairs. This provides strong independent evidence that the assembly is both highly complete and structurally accurate. Together, these results demonstrate the potential of our approach to generate high quality draft genome assemblies from single cells of novel, uncultivated microorganisms.

While our results demonstrate that we can overcome high chimera rates, overall assembly contiguity and genome completeness are ultimately constrained by the uneven amplification of MDA. Even when overall sequencing depths are high, coverage dropouts will result in fragmented assemblies, a consequence of starting from just a single copy of the genome. Overcoming these limitations will require improvements to WGA methods. Encouragingly, several directions show promise. Engineered variants of the phi29 DNA polymerase have been developed, for example, the HotJa phi29 variant which operates at a higher temperature of 40°c demonstrated improved coverage relative to wild-type phi29 (Zhang et al. 2023). Methods such as droplet-based MDA (dMDA), where amplification takes place in droplets containing a single DNA fragment, also report improved coverage uniformity compared to traditional MDA (Hård et al. 2023). Primary template-directed amplification (PTA) is a recently developed method based on MDA that takes a different approach (Gonzalez-Pena et al. 2021). PTA also uses phi29 to amplify template DNA but incorporates exonuclease-resistant terminators during amplification. This reduces the size of amplification products and limits their subsequent re-amplification, converting the reaction from an exponential to a quasilinear process in which the majority of amplicons derive from the primary template. PTA has been shown to substantially improve genome coverage breadth and uniformity compared to conventional MDA (Gonzalez-Pena et al. 2021). However, due to the nature of the approach, PTA amplification products are shorter than traditional MDA, potentially limiting their utility for long-read sequencing. We anticipate that future approaches will simultaneously optimise WGA for both coverage uniformity and amplification fragment length, combined with long-read sequencing and computational developments such as lrSAGA, to substantially expand on what can be achieved from a single cell.

## Methods and Materials

### Bulk PacBio HiFi genome sequencing of *Chlamydomonas reinhardtii*

*C. reinhardtii* strain CCAP 11/32A was purchased from the Culture Collection of Algae and Protozoa (CCAP) and maintained in a 1:1 mixture of EG:JM medium (Euglena gracilis medium and Jaworski’s medium). HMW DNA extraction was performed using Nucleon PhytoPure Kit (Cytiva #RPN8511), with a modified version of the manufacturer protocol. Two snap-frozen Chlamydomonas cell pellets, each containing ∼400M cells and weighing ∼280 mg, were ground together under liquid nitrogen for a total grinding time of 9 minutes. The powder was thoroughly resuspended in Reagent 1 using a 10mm bacterial spreader loop until the mixture appeared completely homogeneous. 40 µL of 100 mg/ml RNase A (Qiagen #19101) was added, and the sample was mixed with a spreader loop again. The sample was then incubated at 37°C for 30 minutes, at which point Reagent 2 and a further 20 µL of RNase were added and again thoroughly mixed with a spreader loop. The incubation at 65°C was extended from the recommended 10 minutes to 20 minutes. After the incubation on ice, 2 mL of chloroform was added (without resin) and the sample was gently mixed on a 3D platform rocker at room temperature for 10 minutes. 300 µL of Phytopure resin was then added and the sample was mixed on the platform rocker for a further 10 minutes. The upper phase from the subsequent 1300g centrifugation was transferred to a new tube and supplemented with an equal volume of 25:24:1 phenol:chloroform:isoamyl alcohol, mixed gently at 4°C for 10 minutes, and spun at 3000g for 10 minutes. The upper phase from this procedure was transferred to another 15 mL Falcon tube and precipitation proceeded as recommended by the manufacturer protocol.

A total of 7 µg of gDNA was manually sheared with the Megaruptor 3 instrument (Diagenode, P/N B06010003) according to the Megaruptor 3 operations manual and underwent SMRTbell Clean up bead (PacBio®, P/N 102-158-300) purification and concentration before undergoing library preparation using the SMRTbell® Prep Kit 3.0 (PacBio®, P/N 102-141-700). The HiFi library was prepared according to the HiFi gDNA protocol version 02 (PacBio®, P/N 102-166-600), and the final library was size selected using the BluePippin system (Sage Science®), with a 0.75% cassette, S1 marker, and an 8 kb cut-off (Sage Science®). The library was quantified using fluorescence (Invitrogen Qubit™ 3.0, P/N Q33216), and the library size was estimated from a smear analysis performed on the FEMTO Pulse® System (Agilent, P/N M5330AA). The final library was purified with DNeasy PowerClean Pro Cleanup Kit (QIAGEN®, P/N12997-50) according to the manufacturer’s instructions. The sequencing loading calculations were completed using the PacBio® SMRT®Link ABC Calculator v12.0.0.177059. The standard Revio primer was annealed to the adapter sequence of the HiFi library, and the library was bound to the sequencing polymerase with the Revio Polymerase Kit (PacBio®, P/N 102-739-100). Calculations for primer and polymerase binding ratios were kept at default values for the library type. The sequencing chemistry used was Revio Sequencing Plate (PacBio®, P/N 102-118-800) and the Instrument Control Software v12.0.4.197734. The library was sequenced on one Revio 25M SMRTcell. The parameters for sequencing were diffusion loading, 24-hour movie, kinetics off, and 250 pM on-plate loading concentration per cell.

HiFi reads were downsampled prior to assembly by selecting reads ≥ 20 kb, resulting in an estimated coverage of 90.5x. *De novo* genome assembly was performed using hifiasm (version 0.20.0) (Cheng et al. 2021) with default parameters. To curate the assembly, we mapped the subsampled reads against the assembly using minimap2 (Li 2021) and discarded contigs with less than 5x coverage and contigs that contained mitochondrial or plastid sequences. Genome completeness was assessed using BUSCO (version 5.7) (Manni et al. 2021) in genome mode with Augustus using the chlorophyta_odb10 dataset. The mitochondrial and plastid genomes were assembled separately using Oatk (version 1.0) (Zhou et al. 2025).

### Cell sorting and whole genome amplification of *Chlamydomonas reinhardtii*

All plates, tips, seals and vessels were UV-C treated for 30 minutes prior to use. Working in a laminar flow hood, 2 µL of sterile 1×PBS was added to each well of a sterile 96-well plate. The plate was held on ice until use. A 1 mL aliquot of Human K562 cells cultured in RPMI-1640 was stained with 50 µL Propidium iodide (Invitrogen), incubated for 5 minutes at room temperature and set aside on ice. Using a S8 spectral FACS instrument (Becton Dickinson), a single K562 cell from a population exhibiting low PI fluorescence was sorted into each well of row H of the plate containing PBS. The plate was sealed, briefly spun down and held on ice. A 1 mL aliquot of *C. reinhardtii* CCAP 11/32A cells was transferred to an Eppendorf tube with care being taken only to collect cells from the middle of the flask avoiding those at the surface or sitting at the bottom. *C. reinhardtii* cells were loaded onto the sorter. The gating (**Fig. S1B**) was set up as follows: The main body of cells was first gated to exclude very small or very large events based on forward and side scatter. From this, a second gate (‘Singlets’) was selected to exclude possible multi-cell events. Finally, singlet events exhibiting strong autofluorescence in the red channel (625 nm emission) with the violet laser (405 nm) excitation were selected as the population from which to sort. Two *C. reinhardtii* cells from the final gate were sorted into each well of row B and a single *C. reinhardtii* cell into each well of rows C, D, E, F & G of the plate containing K562 cells in row H (**Fig. S1A**). The plate was sealed, briefly spun down then immediately transferred to −70°C Storage.

All tips, tubes, plates, seals and Dragonfly Discovery (SPT Labtech) consumables were UV-C treated for 30 minutes prior to use. All manual steps were carried out in a HEPA filtered laminar flow hood. The plate of sorted cells in PBS was thawed on ice then briefly centrifuged. A master mix comprising 198 µL reconstituted Buffer DLB and 18 µL DTT (Repli-g Single Cell Kit, Qiagen) was prepared. Using a Dragonfly Discovery liquid dispenser (SPT Labtech), 1.5 µL of master mix was added to each sample well. The plate was sealed and briefly centrifuged then incubated using a Thermomixer C (Eppendorf) with a 96-well block and a heated lid at 65°C for 10 minutes. Using the Dragonfly Discovery, 1.5 µL Stop Solution (Qiagen) was added to each reaction. The plate was sealed and then incubated at room temperature, protected from light, for 5 minutes. An amplification mastermix was prepared as follows: 495 µl nuclease-free water, 1595 µL Reaction Buffer and 110 µL Polymerase (Repli-g Single Cell Kit, Qiagen). Using the Dragonfly Discovery, 20 µL of amplification mix was added to each reaction. The plate was sealed, briefly centrifuged then incubated at 30°C for 4 hours using a Thermomixer C (Eppendorf) with a 96-well block and a heated lid. Amplified samples were stored at 4°C. A 1:10 dilution of the amplified gDNA samples was prepared with Low-TE. The concentrations of the 1:10 dilutions amplified gDNA samples were assessed using a fluorescence assay (Quant-it HS DNA, Invitrogen) according to the vendor’s standard protocol and measured on a VANTAstar plate reader (BMG).

### Illumina LITE2.0 single-cell genome sequencing of *Chlamydomonas reinhardtii*

Library preparation and sequencing was delivered by the Technical Genomics Group at the Earlham Institute (Norwich, UK) using LITE2, a modified Illumina DNA Prep protocol (#20060059). All liquid handling steps, except where stated, were carried out on Fluent 780 liquid handling robots (Tecan). All magnetic pull-downs were carried out using a Magnum FLX magnet (Alpaqua) except where stated. All incubations and thermocycler steps were performed using an ODTC thermocycler (Inheco). The concentration of the amplified DNA samples was assessed using a fluorescence assay (Quant-iT HS DNA, Invitrogen). For each sample, a maximum of 15 ng of gDNA in 3.75 µL was combined with 2.5 µL Tagmentation mastermix; comprising a 50:50 mix of Bead-Linked Transposomes and Tagmentation Buffer 1 (BLT & TB1, Illumina). The reactions were shaken for 1 minute at 1500 rpm then incubated at 55°C for 15 minutes followed by 10°C for 5 minutes. 1.25 µL of Tagmentation Stop Buffer (TSB, Illumina) was added to each reaction. The reactions were shaken for 1 minute at 1500 rpm then incubated at 37°C for 15 minutes then 10°C for 5 minutes. The bead-bound DNA was pulled down by a magnet and the supernatant discarded. The beads were then washed three times in 12.5 µL Tagmentation Wash Buffer (TWB, Illumina) per reaction. The beads were resuspended in 2.5 µL of Enhancement PCR mix (EPM, Illumina) and 3.75 µL unique dual inde× 10bp primers pairs (IDT) at 2 µM was added. The DNA was amplified by PCR (68°C for 3 minutes, 98°C for 3 minutes, 7 cycles of 98°C for 45 seconds; 62°C for 30 seconds; and 68°C for 2 minutes, 68°C for 1 minute, and 10°C for 5 minutes). The PCR reactions were subjected to a double-sided size selection (0.5X followed by 0.7X) using Illumina Purification Beads (IPB Illumina) and washed with 80% ethanol. The libraries were eluted in Resuspension Buffer (RSB, Illumina) to a final volume of 12 µl. The concentration of the libraries was assessed using a Quant-iT HS DNA assay (Invitrogen). The fragment size distribution of a subset of libraries was determined using a Bioanalyzer HS DNA assay (Agilent) with a smear analysis size range of 200-1000 bp. This information was used to prepare an equimolar pool of the libraries. An aliquot of the pool was manually cleaned up with 0.7X AMPure XP (Beckman Coulter), 80% ethanol and a magnetic tube rack (Invitrogen). The library pool was eluted in EB to a final volume of 50 µL. The library pool was quantified by fluorescence (Qubit HS DNA, Invitrogen), Bioanalyzer HS DNA assay (Agilent) and finally by qPCR using Illumina Quantification assay (KAPA).

The library pools for skim sequencing (N=94) were diluted down to 500 pM in 24 uL volume prior to adding 0.5 of 0.25nM Illumina PhiX control v3 (Illumina FC-110-3001). A P1 300 cycle NextSeq 1000/2000 P1 reagents (300 cycles) kit with standard SBS reagents (Illumina #20050264) was thawed overnight and 24ul of the library pool dilution with PhiX was added to the reagent cartridge before loading on a NextSeq 1000 and sequenced with 150 bp paired end reads. The six libraries that were selected for deep sequencing were cherry picked from the original LITE2 plate, pooled, and sequenced on a NextSeq 1000 P1 reagents (300 cycles) flow cell following the same procedure.

### PacBio HiFi single-cell genome sequencing of *Chlamydomonas reinhardtii*

DNA from the six selected samples was divided into two aliquots. One of each of the aliquots was subjected to S1 nuclease digestion. For S1 nuclease digestion, 2 μg of DNA from each sample was aliquoted into a new tube and the volume made up to 20 μl to a final concentration of 100 ng/μl. A mastermix was prepared as follows: 3 μl S1 nuclease, 90 μl 5X buffer, and 57 μl nuclease-free water (NEB, EN0321) and held on ice. 10 μl of the S1 mastermix was added to each sample aliquot and gently tip mixed. The reactions were incubated at room temperature for 30 minutes. The reactions were stopped by the addition of 3 μl EDTA at 0.5 M and incubation at 70°C for 10 minutes. Samples were cleaned up with 1X AMPureXP and 80% ethanol and eluted in 25 μl Low-TE.

The SMRTbell libraries were prepared from 1 µg per sample according to the HiFi protocol version 03 (PacBio®, P/N 101-853-100) and barcoded with SMRTbell® Barcoded Adapter Plate 3.0 (PacBio®, P/N 102-009-200). The libraries were quantified by fluorescence (Invitrogen Qubit™ 3.0, P/N Q33216), and the size of each library was estimated from a smear analysis performed on the FEMTO Pulse® System (Agilent, P/N M5330AA). The 12 libraries were pooled equimolarly before sequencing. The sequencing loading calculations were completed using the PacBio® SMRT®Link ABC Calculator v13.1.0.221970. The standard Revio Sequencing primer was annealed to the adapter sequence of the HiFi library pool, and the library pool was bound to the sequencing polymerase with the Revio Polymerase Binding Kit (PacBio®, P/N 102-739-100). Calculations for primer and polymerase binding ratios were kept at default values for the library type. The sequencing chemistry used was Revio Sequencing Plate (PacBio®, P/N 102-118-800) and the Instrument Control Software v13.1.0.221972. The library pool was sequenced on two Revio 25M SMRTcells. The sequencing parameters per cell were adaptive loading, 24-hour movies, Kinetics data off, molecule mode, and 250 pM on-plate loading concentration.

### Analysis of *Chlamydomonas reinhardtii* single-cell genome sequencing data

The complexity of Illumina libraries was estimated by counting unique k-mers and plotting them as a function of the number of reads using bbcountunique from the BBTools toolset. Reads were taxonomically classified using Kraken2 (Wood et al. 2019). Reads classified as “Primates” or a lower taxonomic rank were discarded as contamination. To identify putative chimeras, we mapped PacBio HiFi reads against our reference genome assembly using minimap2 (Li 2021). The percentage of chimeric reads was determined as the proportion of primary alignments that had supplementary alignments (SA:Z tag), identified using SAMtools (Danecek et al. 2021). Structural variant calling was performed using Sniffles2 (version 2.5) (Smolka et al. 2024) for each individual PacBio HiFi library with default parameters. Coverage statistics were calculated using QualiMap (version 2.3) (Okonechnikov et al. 2016). Statistics for Loenz plots were calculated using bam-lorenz-coverage (Hoogstrate et al. 2021).

To benchmark assembly algorithms, Illumina short-reads were assembled using SPAdes (version 4.2.0) (Bankevich et al. 2012) with single-cell mode enabled (--sc) and k-mer sizes 21, 33, 55, and 77. Long-reads from the twelve PacBio HiFi libraries were assembled using Flye (version 2.9.6) (Kolmogorov et al. 2019), hifiasm (version 0.25.0) (Cheng et al. 2021), metaFlye (version 2.9.6) (Kolmogorov et al. 2020), hifiasm-meta (version 0.3-r078) (Feng et al. 2022), and metaMDBG (version 1.2) (Benoit et al. 2024), with default parameters. For lrSAGA assemblies, HiFi reads were error corrected using hifiasm (version 0.25.0) (Cheng et al. 2021) and an assembly graph was constructed using MBG (version 1.0.17) (Rautiainen and Marschall 2021) with a k-mer size of 1001 and then processed using lrSAGA. Assembly graphs were visualised using Bandage (version 0.8.1) (Wick et al. 2015). Assemblies were compared against our reference *C. reinhardtii* CCAP 11/32A assembly using QUAST (version 5.3.0) (Gurevich et al. 2013). Consensus quality values (QVs) were estimated using Merqury (version 1.3) (Rhie et al. 2020), comparing k-mers in the bulk PacBio HiFi reads to the MDA assemblies.

### Assembly of PacBio HiFi and Oxford Nanopore long-read sequencing data from MDA amplified *Caenorhabditis* elegans DNA

PacBio HiFi reads were downloaded from the Sequence Read Archive (SRR27483171) from Roberts et al. (2024), which were generated from half of a single individual *C. elegans* Bristol N2 strain. The HiFi reads were error corrected using hifiasm (version 0.25.0) (Cheng et al. 2021) and an assembly graph was constructed using MBG (version 1.0.17) (Rautiainen and Marschall 2021) with a k-mer size of 2001 and then processed using lrSAGA. An assembly was also generated using hifiasm (version 0.25.0) with default parameters for comparison.

ONT reads were downloaded from the Sequence Read Archive (SRR24201716) from Lee et al. (2023), which were generated from a single individual *C. elegans* Bristol N2 strain. The ONT reads were generated from DNA digested by T7 endonuclease for 30 minutes using a R9.4.1 flow cell. ONT reads were error corrected using NECAT (version 0.0.1) (Chen et al. 2021) and an assembly graph was constructed using MBG with a k-mer size of 1001 and then processed using lrSAGA. We also downloaded the published genome assembly (GCA_030248285.1) from Lee et al. (2023). The PacBio and ONT *C. elegans* assemblies were compared to the reference *C. elegans* Bristol N2 genome (GCF_000002985.6) using QUAST (version 5.3.0) (Gurevich et al. 2013).

### PacBio HiFi single-cell sequencing of protist cells isolated from the environment

Surface water (∼1 m) was collected from the River Leam (52.287295, −1.547563), Royal Leamington Spa (UK) in August 2022, supplemented with 2–3 autoclaved barley grains to support heterotrophic/mixotrophic growth via bacteria increase, and incubated for 1 week prior to cell sorting. Single-cell nuclei were stained with 1xSybrGreen for 10 minutes and sorted into 96-well microplates (pre-filled with 5 µL autoclaved/sterile filtered media), with a BD FACSMelody™ Cell Sorter.

WGA was performed using MDA as part of a modified G&T-seq protocol (Macaulay et al. 2015). Dynabeads MyOne Streptavidin C1 (Invitrogen) beads were washed according to the manufacturer’s guidance in a HEPA filtered laminar flow hood and then incubated with 2× binding & wash buffer (10 mM Tris-HCl [pH 7.5], 1 mM EDTA, and 2 M NaCl) and biotinylated oligo-dT primer (IDT; 5′-/BiotinTEG/AAG CAG TGG TAT CAA CGC AGA GTA CTT TTT TTT TTT TTT TTT TTT TTT TTT TTT TVN-3′) at 100 µM for 30 minutes at room temperature on a rotator. The oligo-treated beads were washed four times in 1× binding & wash buffer (5 mM Tris-HCl [pH 7.5], 0.5 mM EDTA, and 1 M NaCl) and then suspended in 1× SuperScript II First Strand Buffer (Invitrogen) supplemented with SUPERaseIn RNase Inhibitor (Invitrogen) to a final concentration of 1 U/µL. The lysate was thawed on ice. 10 µL of prepared oligo-dT beads was added to each well containing 12 µL cell lysate using a Dragonfly Discovery liquid dispenser (SPT Labtech). The lysate plate was sealed and incubated on a ThermoMixer C (Eppendorf) with a heated lid at 21°C for 20 minutes, shaking at 1,000 rpm. Using a Fluent 480 liquid handling robot (Tecan) and a Magnum FLX magnetic separator (Alpaqua), the lysate supernatant was transferred to a new plate and the beads were washed twice in a custom wash buffer (50 mM Tris-HCl [pH 8.3], 75 mM KCl, 3 mM MgCl2, 10 mM DTT, and 0.5% Tween-20). The supernatant from the washes was added to the leftover cell lysate containing the genomic DNA, which was stored at −20°C overnight. The mRNA was not used for this study. The remaining cell lysate was thawed and subjected to a 0.6× vols Ampure XP clean-up with 80% ethanol. The bead-bound gDNA was isothermally amplified for 3 hours at 30°C then 10 minutes at 65°C using a miniaturized (1/5 vols) Repli-g Single-Cell assay (Qiagen). The amplified gDNA was cleaned up with 0.8× vols Ampure XP and 80% ethanol and then eluted in 10 mM Tris-HCl. Note that these samples weren’t treated with S1 nuclease.

The HiFi libraries were prepared from 159-479 ng of MDA DNA, according to the instructions in the low input protocol version 05 (PacBio®, P/N 101-730-400) and barcoded with SMRTbell® Barcoded Adapter Plate 3.0 (PacBio®, P/N 102-009-200). The libraries were quantified by fluorescence (Invitrogen Qubit™ 3.0, P/N Q33216), and the size of each library was estimated from a smear analysis performed on the FEMTO Pulse® System (Agilent, P/N M5330AA). The four libraries were pooled equimolarly before sequencing. The loading calculations for sequencing were completed using the PacBio® SMRT®Link Binding Calculator v12. Sequencing primer 3.2 was annealed to the adapter sequence of the library pool. Binding of the library pool to the sequencing polymerase was completed using Sequel® II Binding Kit 3.2 (PacBio®, P/N 101-842-900). Calculations for primer to template and polymerase to template binding ratios were kept at default values. Sequel® II DNA internal control was spiked into the library pool complex at the standard concentration before sequencing. The sequencing chemistry used was Sequel® II Sequencing Plate 2.0 (PacBio®, P/N 101-820-200) and the Instrument Control Software v12. The library pool was sequenced on one Sequel II SMRT®cell 8M on the Sequel IIe instrument. The parameters for sequencing were adaptive loading (0.85 target and 2-hour maximum loading time), CCS sequencing mode, 30-hour movie, 2-hour pre-extension time, and 104pM on plate loading concentration.

PacBio HiFi reads were error corrected using hifiasm (version 0.25.0) (Cheng et al. 2021) and an assembly graph was constructed using MBG (version 1.0.17) (Rautiainen and Marschall 2021) with a k-mer size of 1001 and then processed using lrSAGA. Haplotypic duplications were identified and removed using Redundans (version 2.0.1) (Pryszcz and Gabaldón 2016). Assemblies were manually curated to remove bacterial co-bionts using a combination of binning with MetaBat2 (version version 2.15) (Kang et al. 2019) and taxonomic classification using CAT (version 5.3) (von Meijenfeldt et al. 2019) and Tiara (version 1.0.1) (Karlicki et al. 2022). Published bulk RNA-seq reads from *Diphylleia rotans* were downloaded from the Sequence Read Archive (SRR5997434) (Brown et al. 2018) and aligned against the *D. rotans* genome assembly using HISAT2 (version 2.2.1) (Kim et al. 2019).

De novo repeat libraries were generated for each assembly using RepeatModeler (version 2.0.5) (Flynn et al. 2020), and assemblies were soft-masked using RepeatMasker (version 4.1.7). Genome annotation of *Naegleria* sp. PL0398 was performed to match our previous short-read based assembly (McGowan et al. 2025). An initial gene set was predicted using GeneMark-EP (Brůna et al. 2020), with hints generated by ProtHint from a protein database of heterolobosean proteomes. The initial gene set was filtered by selecting complete gene models based on full-length sequence alignments against the same protein database using Diamond (version 2.0.15) (Buchfink et al. 2021) in ultra-sensitive mode. The filtered gene set was used as training genes to train Augustus (version 3.5.0) (Stanke and Morgenstern 2005). The final gene set was predicted using Augustus with the hints generated by ProtHint. The *Bodo* genome was annotated using a similar approach, with hints generated from a protein database of Kinetoplastid proteomes.

For *D. rotans*, an initial gene set was generated using GeneMark-ET (Lomsadze et al. 2014) with the spliced alignments from the bulk RNA-seq data. GeneMark-ET predictions that were supported by the RNA-seq data were used as training genes to train Augustus. Additionally, protein hints were generated using ProtHint with a protein database that included complete transcripts from transcriptome assemblies of related Diphyllatea (Galindo et al. 2024) and the proteome of *Mantamonas sphyraenae* (Blaz et al. 2023). The final gene set was predicted using Augustus, supported by the hints generated from RNA-Seq data and ProtHint.

For *C. triciliatum*, an initial gene set was predicted using GeneMark-EP, with hints generated by ProtHint from the same protein database used above for *D. rotans* but also included the final gene set of *D. rotans*. GeneMark-EP predictions that were supported by ProtHint hints were used as training genes to train Augustus. The final gene set was predicted using Augustus with hints from ProtHint. Genome completeness was estimated for each assembly using BUSCO in protein mode with the eukaryota_odb10 dataset for *C. triciliatum*, *D. rotans*, and *Naegleria* sp., or the euglenozoa_odb10 dataset for *Bodo* sp.

### SSU rRNA phylogenetics

SSU 18S rRNA were annotated using barrnap (version 0.9) (https://github.com/tseemann/barrnap) with “kingdom” set to “euk”. Multiple sequence alignment of *C. triciliatum* and *D. rotans* and related Diphyllatea sequences was performed using MAFFT (version 7.520) with the L-INS-i algorithm (Katoh and Standley 2013). Maximum likelihood phylogenetic reconstruction was performed using IQ-TREE (version 2.3.6) (Minh et al. 2020), under the TPM2u+F+I+R2 model which was the best fitting model determined by ModelFinder (Kalyaanamoorthy et al. 2017). Support was assessed with 100 non-parametric bootstrap replicates.

## Data Availability

Bulk and single-cell sequencing data and the reference genome assembly for *Chlamydomonas reinhardtii* CCAP 11/32A was deposited on the European Nucleotide Archive under BioProject PRJEB85699. The sequencing data and assemblies for the *Bodo*, *Collodictyon*, *Diphyllelia*, and *Naegleria* samples were deposited under BioProject PRJEB111600. Additional supporting data have been deposited on Zenodo (<u>10.5281/zenodo.18268087</u>). lrSAGA is available at https://github.com/jamiemcg/lrSAGA.

## Acknowledgements

We would like to acknowledge the members of the Technical Genomics and Core Bioinformatics groups at the Earlham Institute, and note the specific contributions of Chris Watkins, Kendall Baker, and Neil Shearer. We thank Andrew Goldson of the Single-Cell and Spatial Analysis Group at the Earlham Institute for cell sorting and advice on cytometry data and image analysis. We also thank Charlie Nightingale and George Basnet for assisting with cell culture and collection. We acknowledge the work delivered via the Laboratory Managers and Research Computing Group at the Earlham Institute who manages and delivers High-Performance Computing.

---

**Supplementary Figure 1.**
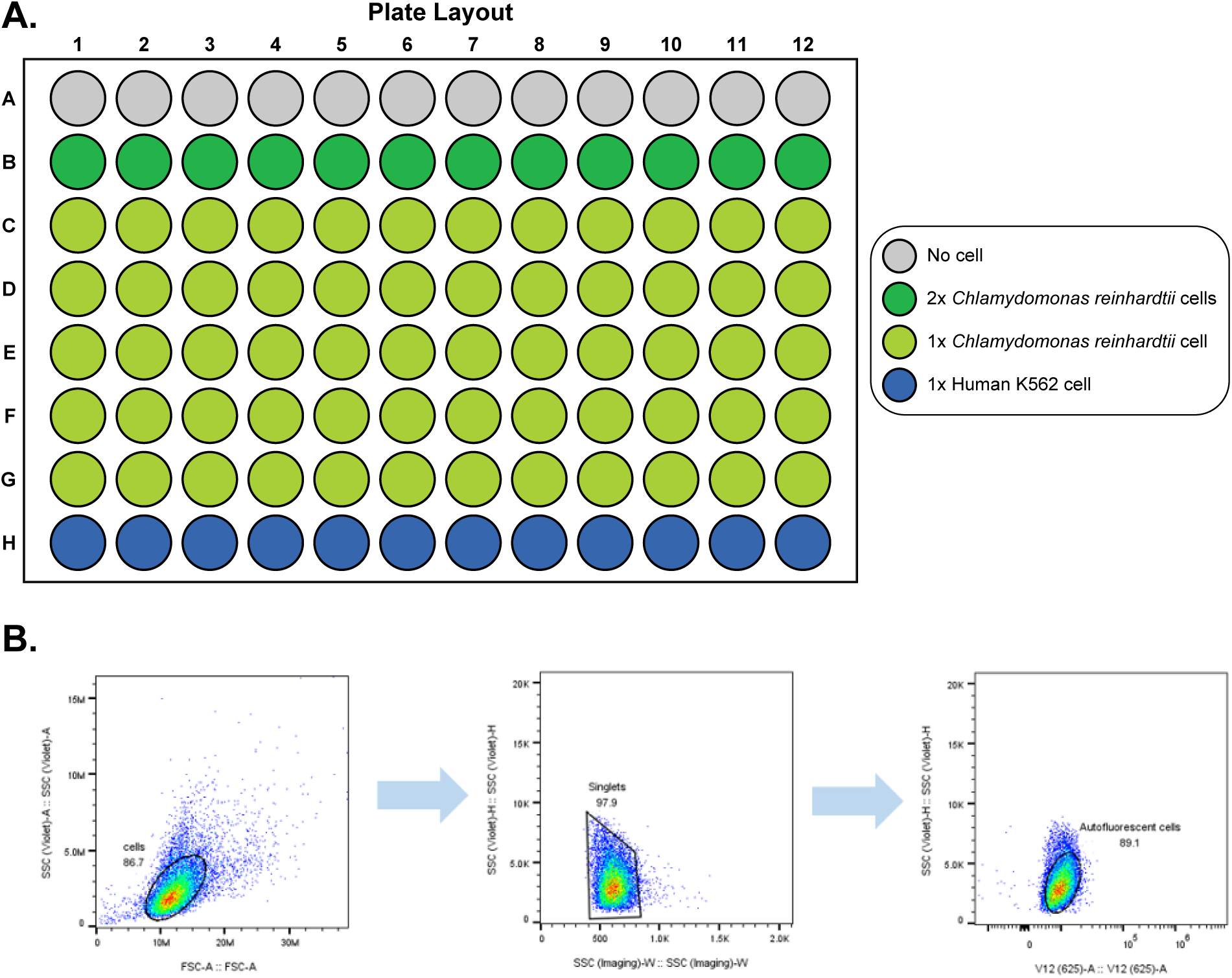
**(A)** 96-well plate layout. No cells were sorted into row A, two cells of *C. reinhardtii* were sorted into each well of row B, a single-cell of *C. reinhardtii* CCAP 11/32A was sorted into each well of rows C to G, and a single Human K562 cell was sorted into the wells of row H. (**B**) Representative FACS gating strategy to sort *C. reinhardtii* cells. The main body of cells was first gated to exclude very small or very large events based on forward and side scatter. From this, a second gate (‘Singlets’) was selected to exclude possible multi-cell events. Finally, singlet events exhibiting strong autofluorescence in the red channel (625 nm emission) with the violet laser (405 nm) excitation were selected as the population from which to sort.

**Supplementary Figure 2.**
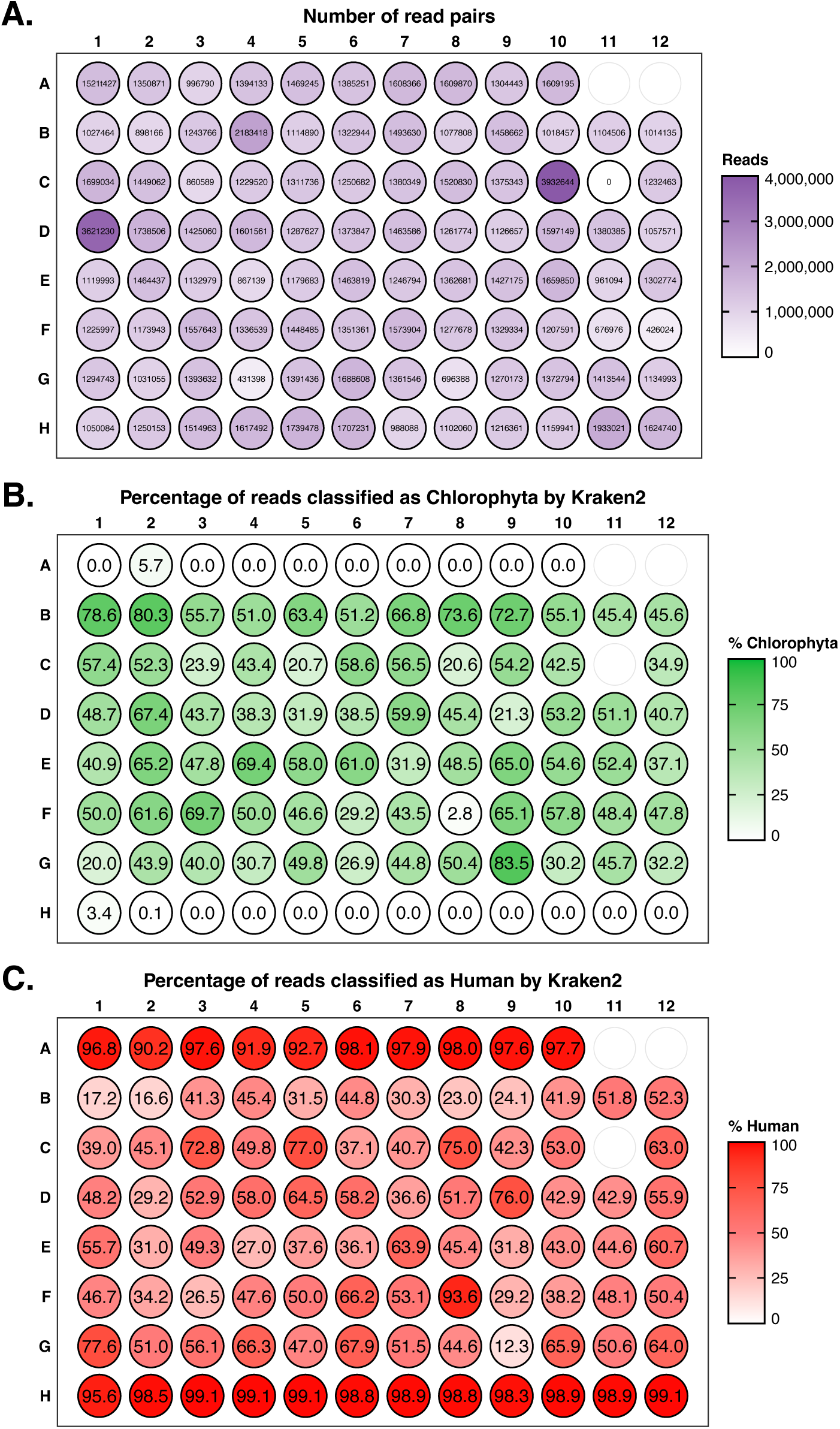
Low-coverage Illumina skim-sequencing of 96-well plate of MDA amplified DNA showing the number of read-pairs recovered per well (**A**), the percentage of reads classified as Chlorophyta by Kraken2 (**B**), and the percentage of reads classified as Human by Kraken2 (**C**). Wells A11 and A12 were not sequenced.

**Supplementary Figure 3.**
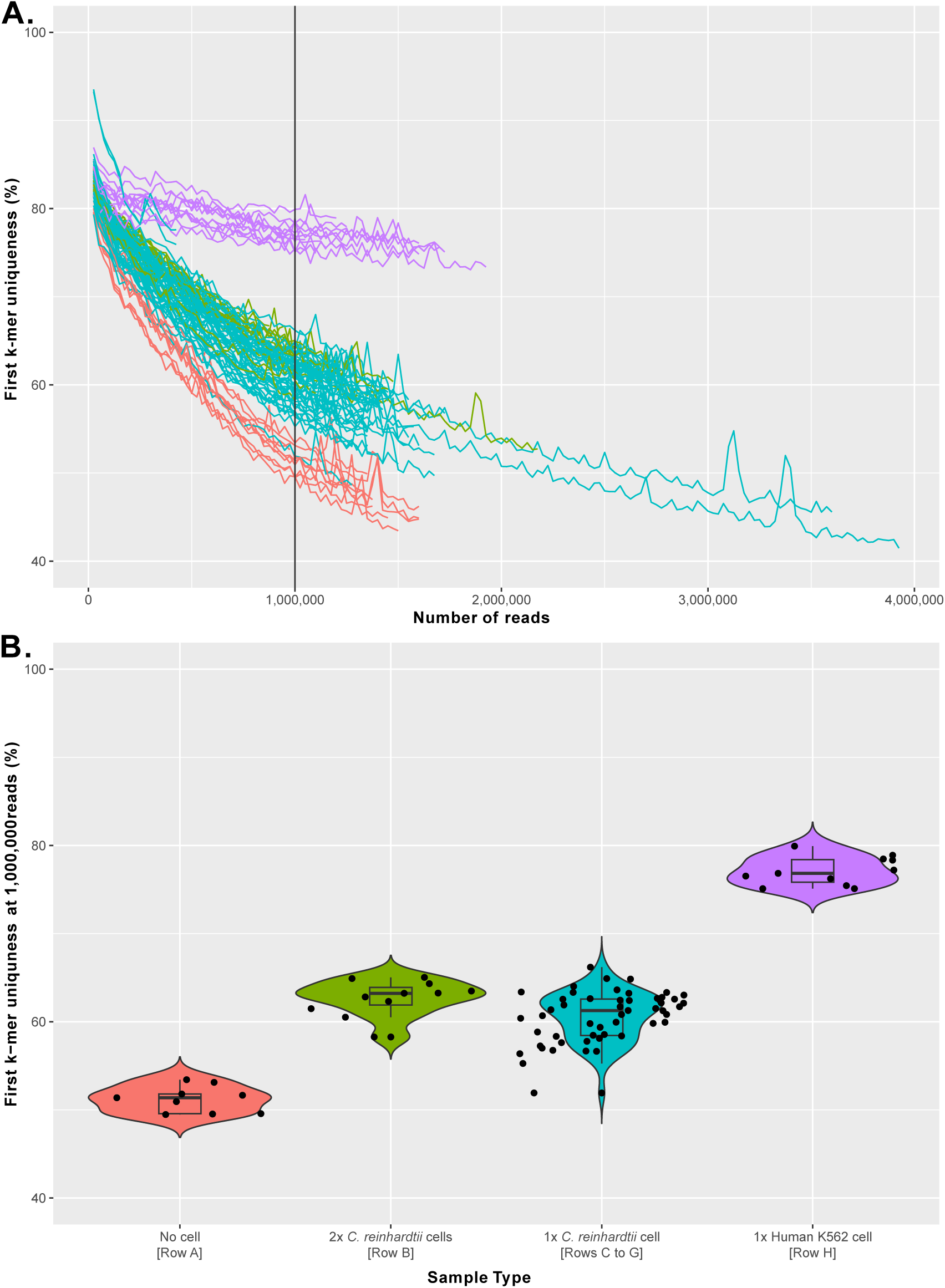
Estimates of library diversity of Illumina skim-seq libraries based on k-mer uniqueness. K-mer uniqueness plotted as a function of the number of reads, with 1,000,000 reads indicated by a vertical bar (**A**). Violin plots show k-mer uniqueness values at 1,000,000 reads with samples grouped by type (no cell, two *C. reinhardtii* cells, a single *C. reinhardtii* cells, or a single Human control cell) (**B**).

**Supplementary Figure 4.**
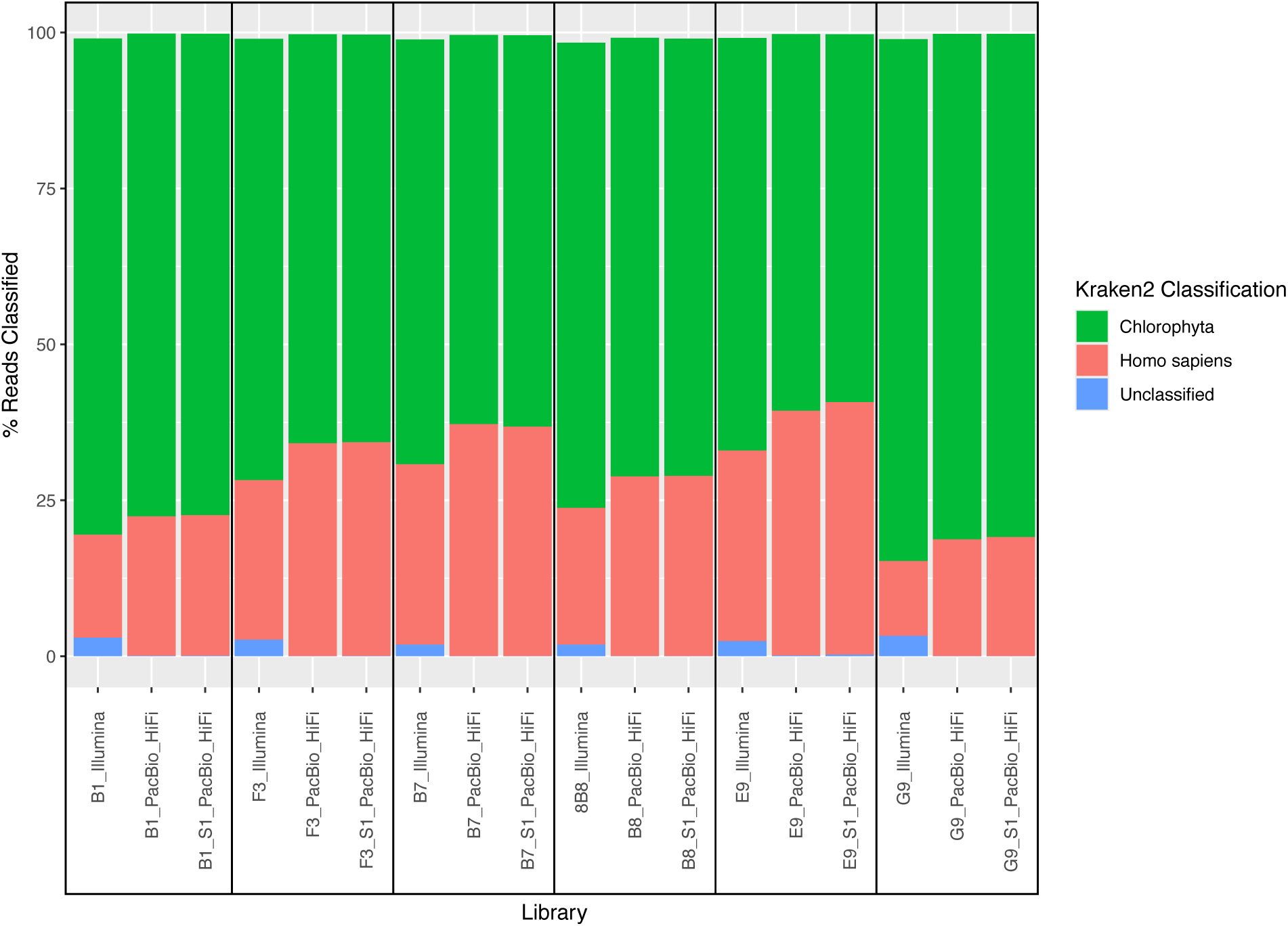
Kraken2 estimates of human contamination for Illumina and PacBio HiFi libraries.

**Supplementary Figure 5.**
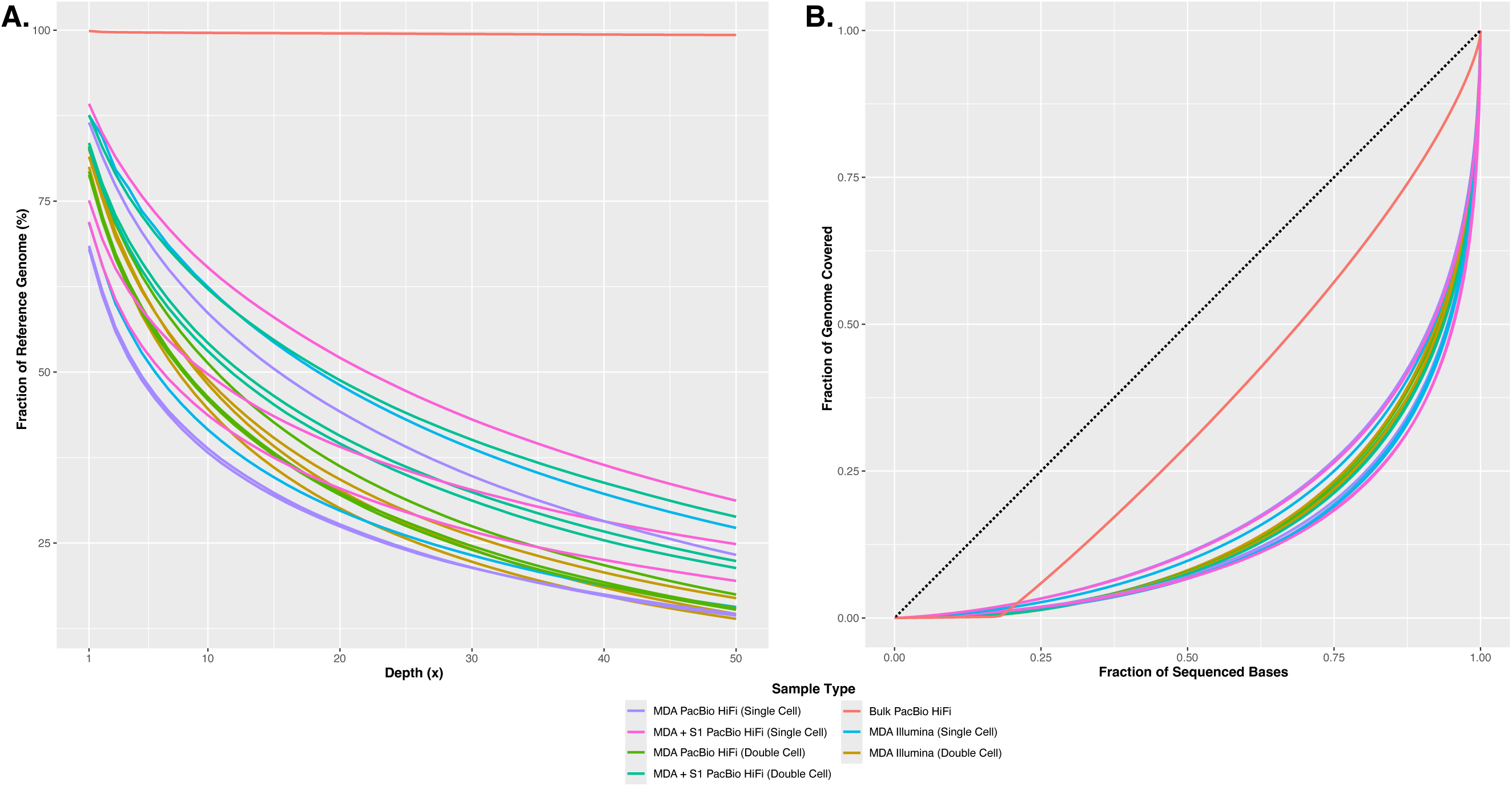
Coverage statistics of *C. reinhardtii* CCAP 11/32A PacBio HiFi and Illumina libraries. Samples are coloured to indicate sample type (bulk, single cells, or double cells), sequencing platform (Illumina or PacBio HiFi), and if samples were digested with S1 nuclease or not (MDA or MDA + S1). (**A**) Genome fraction coverage plot computed using QualiMap showing the fraction of the reference CCAP 11/32A genome that was recovered at each depth up to 50x. (**B**) Lorenz plot illustrating coverage uniformity computed using bam-lorenz-coverage. The dashed line shows theoretical uniform coverage. Note that the high copy number of the plastid genome skews uniformity estimates, most noticeable in the Bulk PacBio HiFi library. Although the plastid genome represents approximately 0.18% of the total reference assembly length at 206 kb, it accounts for approximately 16% of bulk sequencing reads.

**Supplementary Figure 6.**
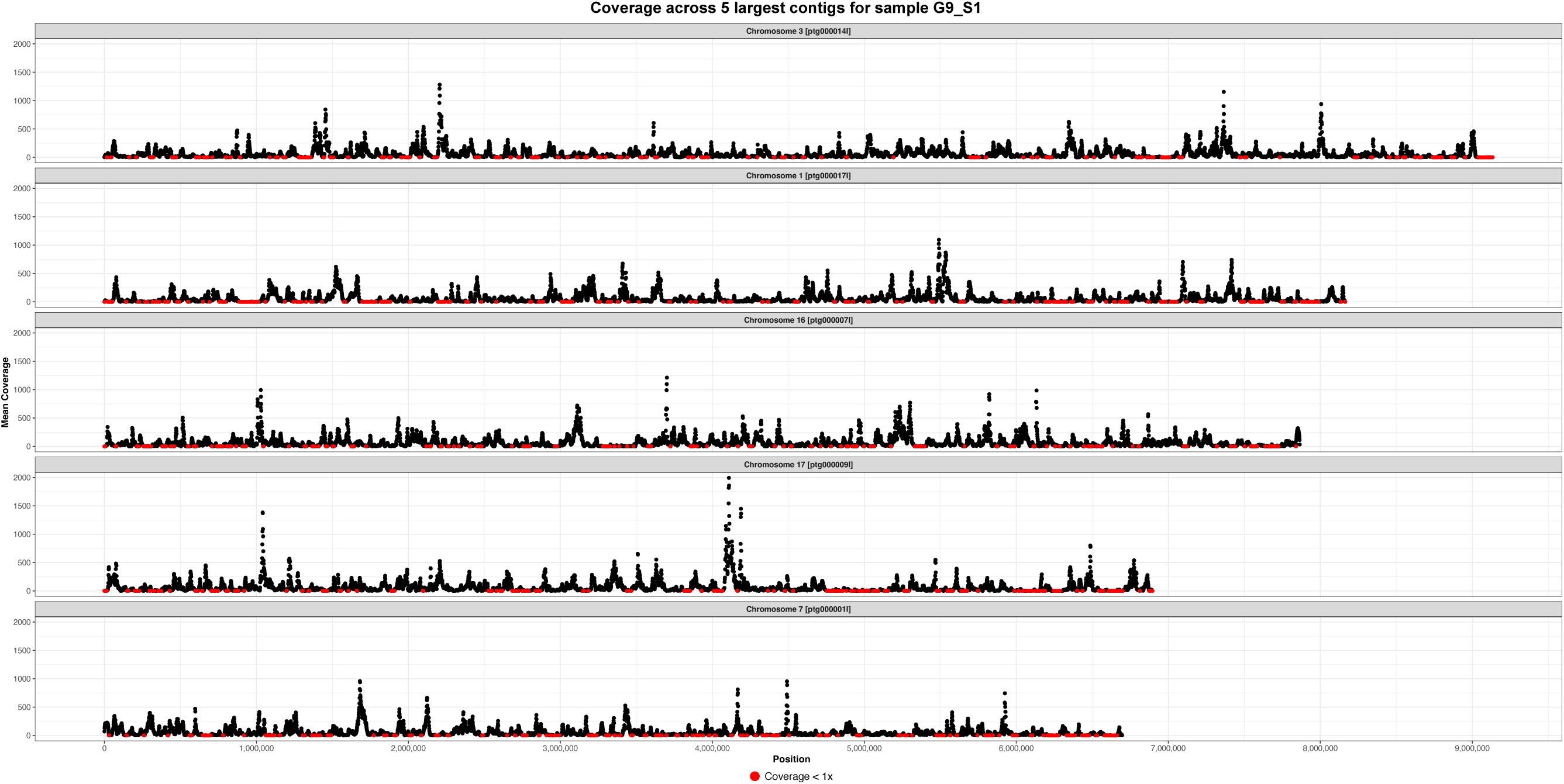
Coverage plots showing uneven coverage distribution of PacBio HiFi MDA library for sample G9_S1 across the five largest contigs from the reference *C. reinhardtii* CCAP 11/32A genome assembly, corresponding to chromosomes 3, 1, 16, 17, and 7. Coverage dropouts (coverage < 1x) are indicated with red dots.

**Supplementary Figure 7.**
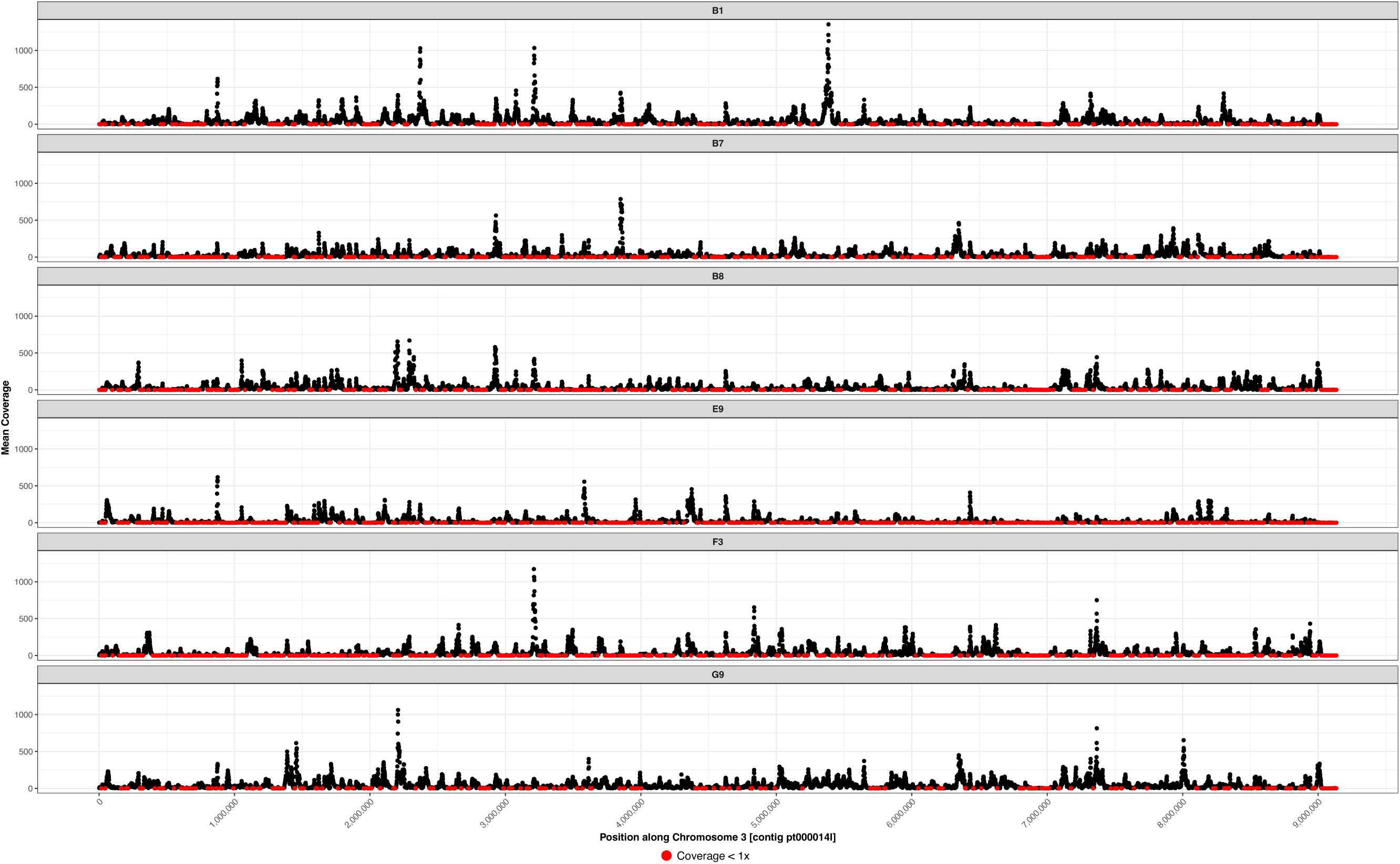
Coverage plots showing uneven coverage distribution from six PacBio HiFi MDA libraries across the largest contig from the reference *C. reinhardtii* CCAP 11/32A genome assembly, corresponding to chromosome 3. Coverage dropouts (coverage < 1x) are indicated with red dots.

**Supplementary Figure 8.**
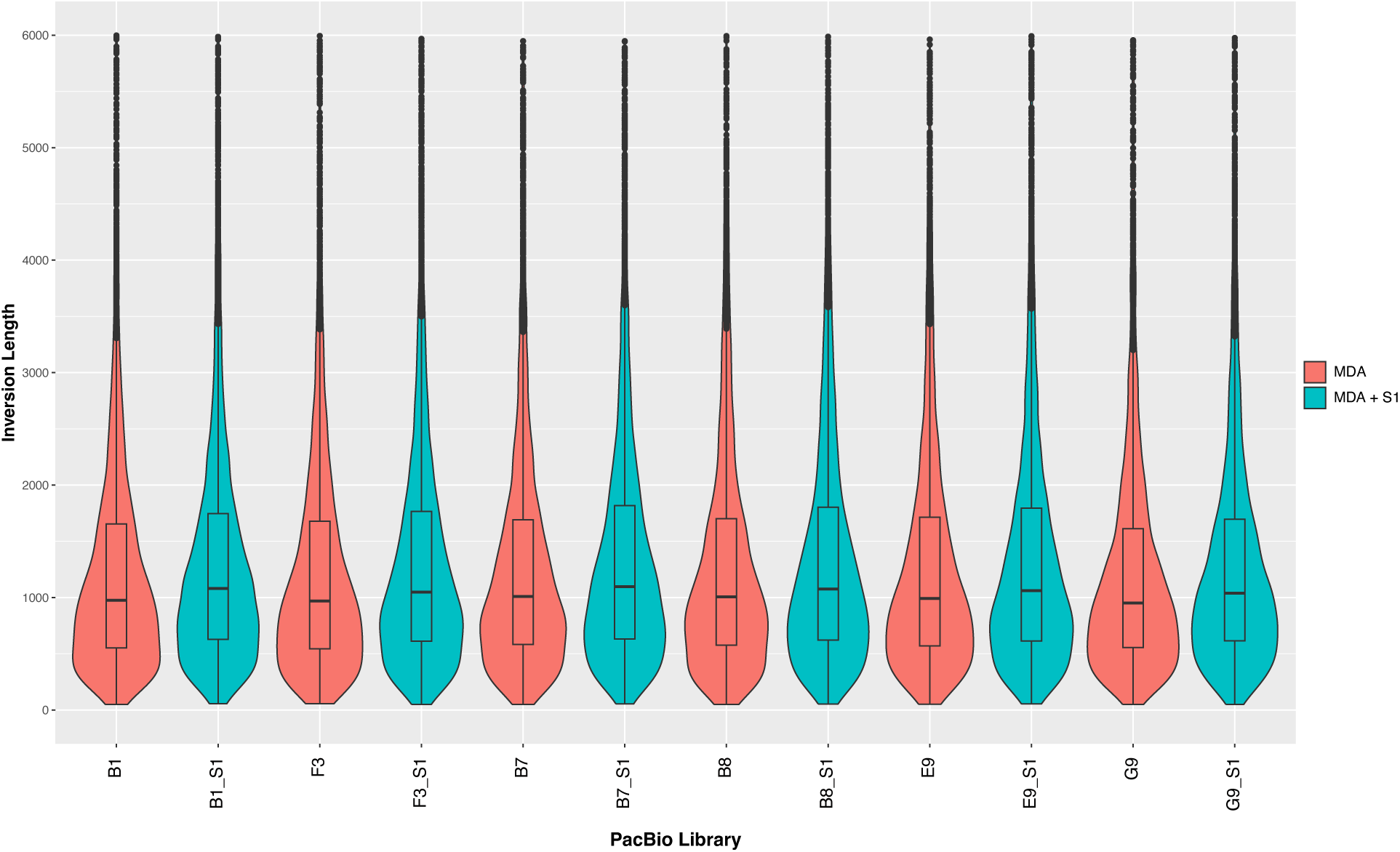
Violin plots showing the distribution of inversion lengths identified by Sniffles2 based on aligning each of the PacBio HiFi MDA libraries against the reference *C. reinhardtii* CCAP 11/32A genome assembly.

**Supplementary Figure 9.**
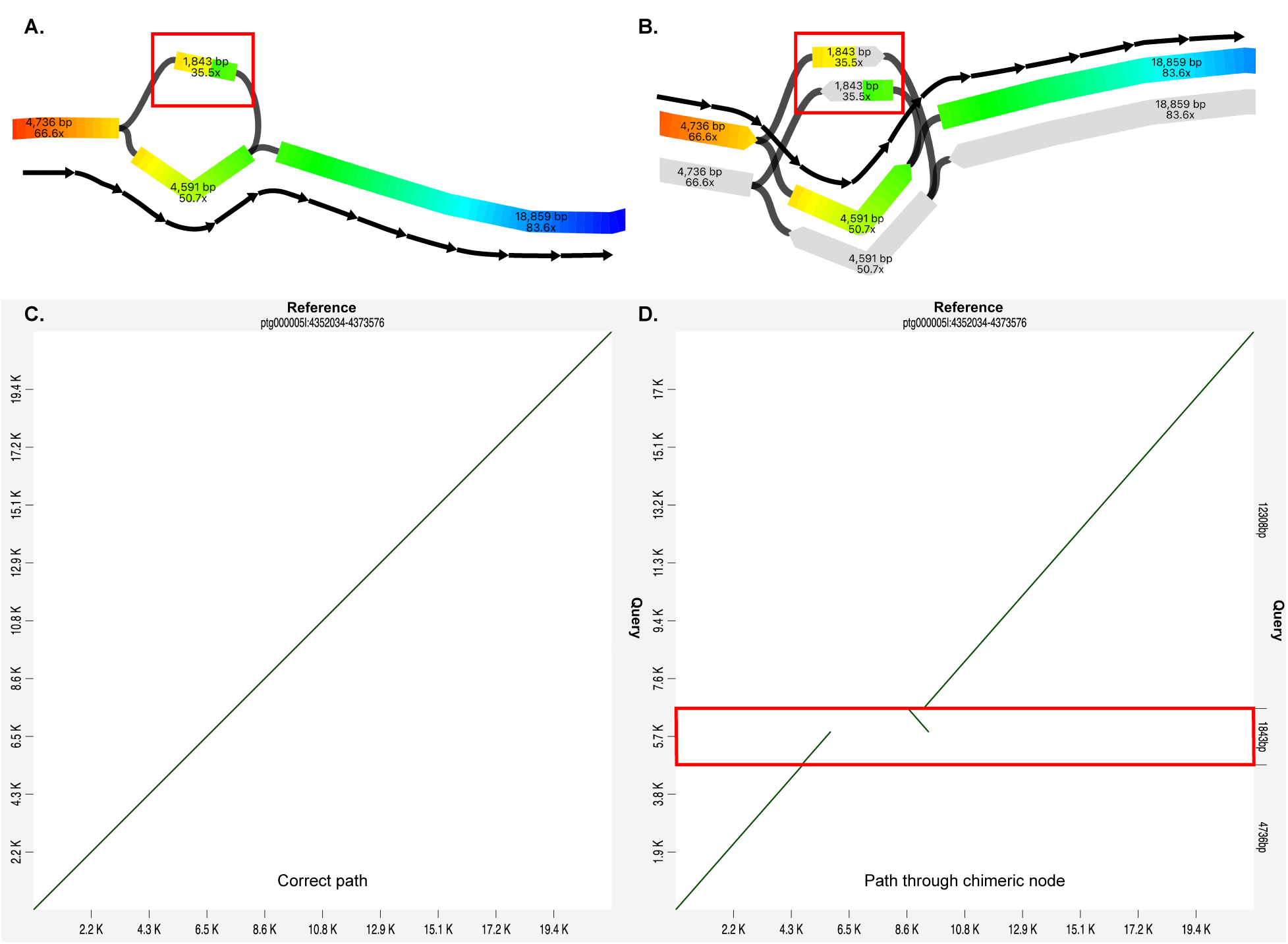
Example of a chimera detected in an assembly graph. (**A**) Bandage visualisation of an assembly graph segment interrupted by an inverted chimera (highlighted by a red box). The correct path through the graph is indicated by black arrows. Nodes are coloured according to their positions in a BLAST search using the correct sequence from the reference assembly a query. A sharp change in colour is visible in the chimeric node, indicating a structural error relative to the reference genome. The chimeric node interrupts an otherwise linear path through the assembly graph (**B**) Bandage visualisation of the same assembly graph segment rendered in “Double node style” where each node and its reverse complement strand are rendered separately. This shows that the chimeric node overlaps with one neighbour in the forward orientation and to another in its reverse complement orientation, looping back in the wrong direction causing an inversion. (**C**) Alignment of the correct path against the reference genome demonstrating a continuous linear alignment. (**D**) Alignment of the path through the chimeric node against the reference genome showing a non-linear alignment interrupted by an inverted chimera.

**Supplementary Figure 10.**
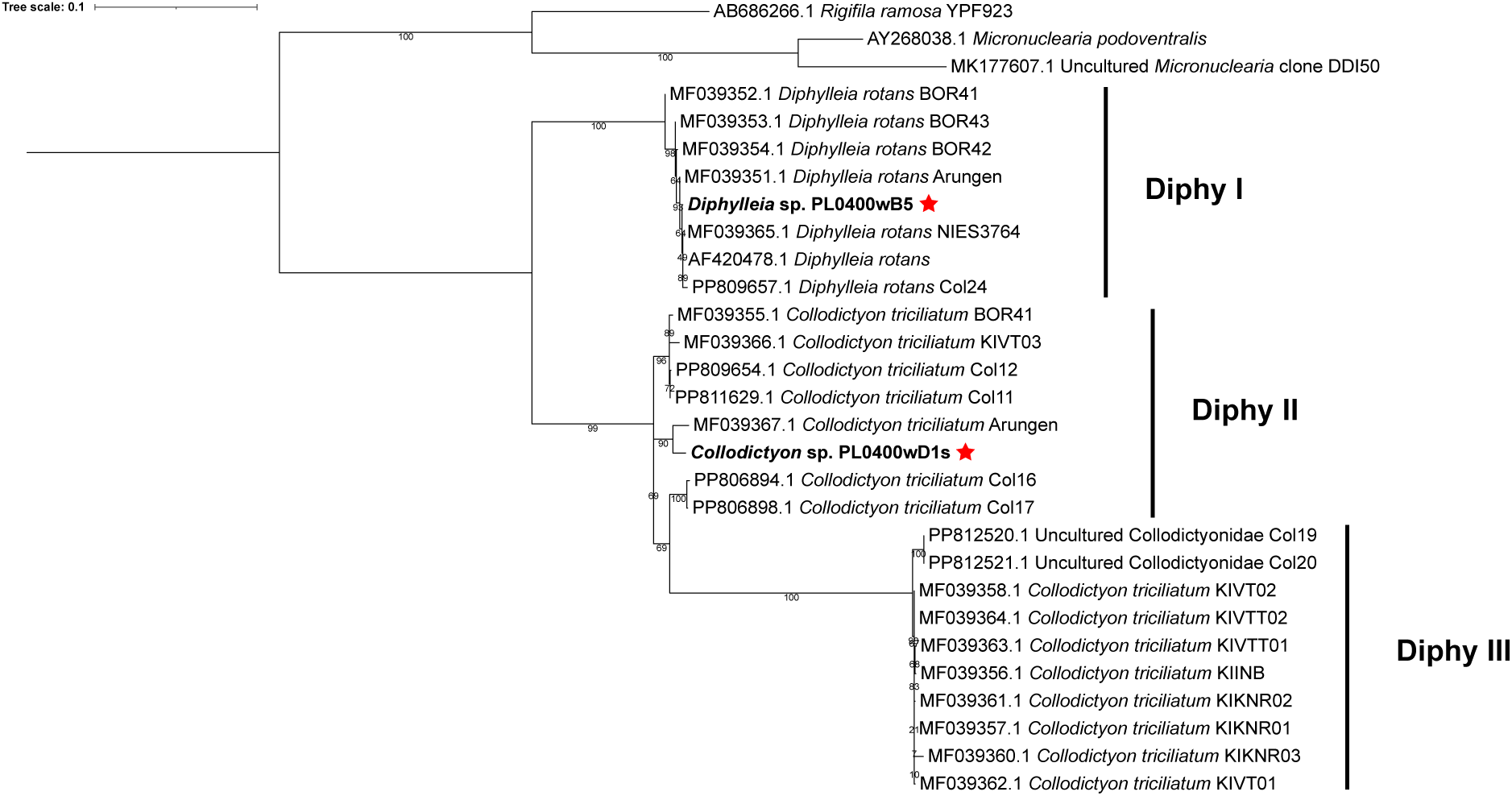
Maximum-likelihood phylogenetic analysis of SSU rRNA genes from Diphyllatea under the TPM2u+F+I+R2 model. Sequences from the two single-cell genome assemblies are indicated in bold with a red star. Branch support values were computed using 100 non-parametric bootstraps.

